# MALDI Deamidation Score (MDS): A Fast and Flexible Method for Assessing Deamidation in ZooMS Data and Its Application to the Denisova Cave Bone Assemblage

**DOI:** 10.1101/2025.08.26.671509

**Authors:** Fei Yang, Ismael Rodríguez Palomo, Bharath Anila Bhuvanendran Nair, Samantha Brown

## Abstract

Estimating deamidation from MALDI-TOF MS spectra of bones has most frequently been achieved using the q2e method due to its high-throughput capacity and ease of use. Despite this accessibility, q2e is only capable of estimating deamidation at the peptide level on a fixed peptide list and does not generate a sample-wide summary. The introduction of the Parchment Glutamine Index (PQI) presented an alternative method for deamidation estimation. Initially designed for a large ZooMS dataset of parchment, it utilises weighted least squares and a linear mixed-effects model (LME) to generate deamidation estimation on both the peptide and sample level. To address the limitations of q2e and expand the applicability of PQI to a wider range of archaeological tissues and MALDI-derived data (such as ZooMS data on bone collagen), we developed the MALDI Deamidation Score (MDS), an iteration of the PQI method optimized for handling large-scale datasets. Compared to PQI, MDS is more streamlined for the analysis of multi-species data with customisable peptide lists, offering a dramatically decreased processing time while being able to normalise the peak intensity for higher accuracy. Through a case study on the published Denisova Cave ZooMS assemblage, we demonstrate that different peptides exhibit varying deamidation patterns over time, making the use of a single peptide to represent overall deamidation potentially biased. Such information is invaluable for investigating key questions such as protein preservation and site formation processes, especially when contextualized with other lines of archaeological evidence.

## Introduction

### Review on application of deamidation in ZooMS

Zooarchaeology by Mass Spectrometry (ZooMS) is a widely used palaeoproteomic technique for identifying the taxonomic origin of collagen-rich archaeological, historical, and palaeontological materials, like bone, dentine, ivory, leather, and manuscripts, by analysing their peptide mass fingerprints and comparing the resulting spectra with established reference libraries (Brandt et al., 2020; Buckley et al., 2009; Coutu et al., 2016; Fiddyment et al., 2015; Richter et al., 2022). Beyond taxonomic identification, researchers have increasingly sought to extract diagenetic information from ZooMS spectra, most notably through the analysis of glutamine deamidation (Antonosyan et al., 2024; Bray et al., 2023; Pal Chowdhury et al., 2019; Peters et al., 2023; Welker et al., 2016), a spontaneous post-translational modification that accumulates over time, known to be influenced by multiple burial conditions, including pH, temperature, or humidity (Scotchler and Robinson, 1974).

Deamidation is the process through which glutamine and asparagine convert to glutamic acid and aspartic acid, respectively. There are two different mechanisms that can induce spontaneous deamidation, the first is through direct hydrolysis of the side chain and the second is through the formation of a cyclical intermediate (Catak et al., 2009; Joshi et al., 2005; Riggs et al., 2019). The process through which asparagine converts to aspartic acid has been well studied, as it is much more sensitive to deamidation than glutamine, and therefore can be observed under shorter timescales (Daniel et al., 1996; Robinson et al., 2004; Terwilliger and Clarke, 1981; Wilson et al., 2012). From an archaeological and palaeontological perspective, glutamine is therefore a more suitable measure of protein diagenesis and has now been routinely observed in bones and teeth dating back over millions of years (Cappellini et al., 2019; Madupe et al., 2025). Given that taphonomy has the potential to influence the rate of deamidation, there has long been a proposition that it may be used as a molecular clock (Robinson et al., 1970) with several attempts to understand its mechanisms and kinetics (see for example (Catak et al., 2009; Hao et al., 2017; Ramsøe et al., 2021; Riggs et al., 2019; Robinson and Rudd, 1974).

The development of ZooMS enabled the practical application of fast and low-cost deamidation analysis for fossil specimens and artefacts, which also required taxonomic identification. This advancement was further bolstered by two methodological advancements. The first was the development of the non-destructive ammonium bicarbonate (AmBic) extraction protocol for type-I collagen (COL1). This protocol was specifically designed to minimize lab-induced deamidation brought on through acid-based demineralisation (van Doorn et al., 2011). The second was the introduction of an isotopic envelope deconvolution method for estimating deamidation levels on a fixed series of peptides from ZooMS spectra (Wilson et al., 2012), known as the ‘q2e method’. This method employs a genetic algorithm to estimate the abundance of the deamidated and non-deamidated peptide counterparts in the mixed isotopic envelope. Within the framework of the q2e method, two collagen peptides - COL1α1 508-519 (*m/z* 1105.58, GVQGPPGPAGPR) and COL1α1 435-453 (*m/z* 1706.77, DGEAGAQGPPGPAGPAGER) - are the most consistently detected in ZooMS spectra and are broadly conserved across terrestrial mammals common in the archaeological record. Therefore, these peptides are the best candidates for deamidation estimation on large faunal assemblage with diverse species to minimize amino acid sequence induced bias (Robinson and Robinson, 2001). However, empirical data show that COL1α1 435-453 is only present in about half of the spectra (Welker et al., 2017), making COL1α1 508-519 the primary marker in most studies focused on ZooMS analyses of fossil bone assemblages.

Despite early research indicating a correlation between the deamidation levels of COL1α1 508-519 and the thermal age, exceptions due to various taphonomic factors have frequently been observed (van Doorn et al., 2012; Wilson et al., 2012). While thermal age tries to set an equivalent thermal history by encompassing multiple environmental factors on an equivalent or normalised time scale, it still misses most of the critical factors and their overall dominance and locality. This complicates the correlation between deamidation levels and chronological or thermal age, especially when samples from diverse regional, environmental and depositional settings are considered, or when there are large intra-site variations across time that are not considered by the thermal age (Pal Chowdhury and Buckley, 2022; Schroeter and Cleland, 2016). To understand the underlying mechanisms that may induce or accelerate deamidation in bone, numerous controlled experiments and empirical studies on archaeological material have been conducted which attempt to explore potential factors influencing deamidation beyond time, including sampling point, bone type, taphonomy and lab protocols for collagen extraction (Ásmundsdóttir et al., 2024; Mylopotamitaki et al., 2023; Pal Chowdhury et al., 2019; Simpson et al., 2016; Welker et al., 2017). Together, they demonstrate the complexity of measuring protein diagenesis even within a single site as the deamidation of consecutive layers does not always follow a linear pattern. These findings, nevertheless, have not dissuaded recent ZooMS studies from continuing to publish deamidation data on COL1α1 508-519 using the q2e method, either as individual measurements or in an attempt to explain observed patterns in stratigraphy or biomolecular preservation (Antonosyan et al., 2024; Brown et al., 2021b; Sinet-Mathiot et al., 2019; Xia et al., 2024). Deamidation of glutamine remains the only post-mortem post-translational modification studied in ZooMS data and is frequently relied upon to demonstrate endogeneity and antiquity of specimens despite the complications with its use. So far, a limited number of ZooMS studies have successfully used deamidation rates to indicate stratigraphic integrity and sample’s authenticity (for example Martisius et al., 2020; Xia et al., 2024).

As an alternative to the q2e method, a new method called Parchment Glutamine Index (PQI) was developed for assessing parchment quality via a sample-level deamidation value and to explore patterns of parchment production during the Medieval period (Nair et al., 2023). To date there has been no publication on the application of PQI on other protein-based archaeological material.

### From q2e to PQI and MDS

ZooMS peptide mass fingerprint spectrum is generated using Matrix-Assisted Laser Desorption/Ionisation (MALDI-TOF) mass spectrometry, a rapid yet low-resolution method of measuring proteins and peptides (Buckley et al., 2010, 2009). While MALDI-TOF MS may limit precise quantification of post-translational modifications, the rapidly growing volume of publicly available ZooMS data offers a valuable source for metadata analysis. These datasets can help reveal diachronic patterns from single archaeological sites or even large-scale geographic areas. The q2e method employs a genetic algorithm to estimate deamidation levels by stochastically generating a series of candidate values and selecting the one that best fits the isotopic envelope of a glutamine-containing peptide (Wilson et al., 2012). Despite its high-throughput nature and widespread use in the field, the q2e method has several limitations. First, although the q2e method generates peptide-level deamidation values, it does not estimate an overall sample-level deamidation value based on multiple peptides. Most applications focus on a single peptide, COL1α1 508-519, as it is widely observed in MALDI-TOF spectra and the peptide sequence remains constant across multiple species. Van Doorn and colleagues (2012) proposed a composite deamidation value based upon the average of different peptides, weighted by its logarithmic correlation with the peptide COL1α1 508-519. This again places more weight on this ubiquitous peptide without any consideration on the variation in peptide-specific deamidation kinetics. Moreover, values derived from sample triplicates are often averaged in a simplistic manner, without considering differences in spectral quality among the replicates. Third, the peptide list for the q2e is not easily configurable for the users to select different sets of peptides depending on the archaeological context. Fourth, it uses a stochastic and non-deterministic genetic algorithm optimization to estimate the deamidation, while the deconvolution can in fact be formalised as a linear least squares optimization problem that can be solved with a mathematical expression (Nair et al., 2023).

By contrast, the Parchment Glutamine Index (PQI) (Nair et al., 2023) uses weighted least-squares (WLS) to estimate deamidation at the peptide level and a linear mixed-effects model (LME) to combine the peptides and replicates estimates on a sample level index, based on a user-defined list of peptides. After spectral preprocessing, each peptide’s deamidation is fitted by weighted least squares (WLS); replicates with higher signal-to-noise ratio (SNR) receive higher weights, and peptides shown to be more responsive to deamidation receive greater influence in the overall sample score. The peptide estimates for each replicate are then integrated and in turn weighted by their WLS fitting residuals, to yield a single PQI value representing sample level deamidation.

To assess this new method’s suitability to the analysis of bone, we adapted the PQI statistical framework to develop the **MALDI Deamidation Score (MDS)** - a streamlined pipeline with WLS-LME architecture tailored for large ZooMS MALDI-TOF datasets. MDS retains the flexibility of PQI while offering improved usability and applicability to diverse peptide sets. We evaluate its performance against the q2e method and apply it to the Denisovan dataset (Brown et al., 2021c), which is currently the largest open-access multi-layer ZooMS dataset from a single archaeological site. This application assesses the potential of MDS to improve deamidation estimation accuracy and to refine our understanding of collagen preservation in archaeological or palaeontological contexts. For clarity, in the following text, peptide-level deamidation estimates generated by MDS are referred to as **Weighted Least Squares q2e (WLS-q2e)** and are subsequently integrated into an MDS score representing sample-level deamidation, whereas deamidation estimates based on pep2 (*m/z* 1105) using the original q2e method developed by Wilson et al. (2012) are referred to as **Genetic Algorithm q2e (GA-q2e)**. In MDS, higher values indicate higher levels of deamidation.

## Material

Denisova Cave, located in the Altai region of Russian Siberia, has yielded valuable ZooMS data collected from its three chambers (Brown et al., 2022, 2021c, 2016). The deamidation analysis (Brown et al., 2022, 2021c, 2016) specifically focused on samples from the East Chamber, where the majority of samples from the site had been analysed using ZooMS. This dataset spans a long stratigraphic sequence, encompassing seven layers over approximately 240,000 years (Douka et al., 2019; Jacobs et al., 2019). Such a comprehensive time range makes it particularly suitable for assessing collagen preservation changes over extended periods and across diverse depositional environments. In total, over 8,000 bone fragments from Denisova Cave were analysed using ZooMS (Brown et al., 2022, 2021c, 2016). Two different extraction protocols were used across the entire dataset, an acid-based protocol and the AmBic protocol developed specifically for the analysis of deamidation (van Doorn et al., 2011). Brown and colleagues (2021) selected 2459 samples for inclusion in their deamidation study. These were chosen following four criteria; i) specimens were excavated from the East Chamber, to give some stratigraphic control, ii) collagen was extracted following a previously published AmBic protocol (van Doorn et al., 2011; Welker et al., 2017; (Brown et al., 2020), iii) each specimen produced three replicate MALDI-TOF MS spectra, iv) each specimen could be taxonomically identified using ZooMS. Spectra were acquired at the Department of Archaeology, Max Planck Institute for the Science of Human History (MPI-SHH) in Jena, Germany. Peptides are reported following standard conventions (Brown et al., 2021a). A total of 2459 samples, each with three replicates, would have resulted in an expected dataset of 7377 spectra; however, only 7358 spectra were retrievable from the online database.

For the present analysis, we focused on the following nine taxonomic groups identified in Brown et al. (2021): *Bison/Yak*, *Capra*, *Ovis*, *Rhinocerotidae*, *Equidae*, *Cervidae/Gazella/Saiga*, *Ursidae*, *Canidae*, and *Crocuta/Panthera*. These nine groups comprise the majority of terrestrial mammalian specimens in the dataset, including both herbivores and carnivores. Moreover, they share several peptides containing at least one glutamine residue, making them well-suited for evaluating the performance of our novel MDS method across multiple taxa.

## Method

### Selection of peptides

We selected seven peptides for the MDS analysis of the nine major taxa groups included in this study (Table 1). Those peptides contain at least one glutamine (with their potential hydroxylated versions), and share the same positions by all nine taxa groups. We focus mainly on those peptides which are visible in ZooMS spectra and are already included in routine taxonomic identifications as they tend to ionise and be detected well by MALDI-TOF MS (COL1α1 508-519, COL1α2 757-789, COL1α2 757-789’, COL1α2 978-990, and COL1α2 978-990’). Two additional peptides, which are not routinely included in ZooMS analysis, were also chosen as they were visible across ZooMS spectra and each contained one glutamine (COL1α1 435-453’ and COL1α1 781-789). The peptide list for *Bison/Yak* is shown here as an example on Table 1, and the full list including all nine taxa is available with the code. See Data and code availability.

**Table 1:**
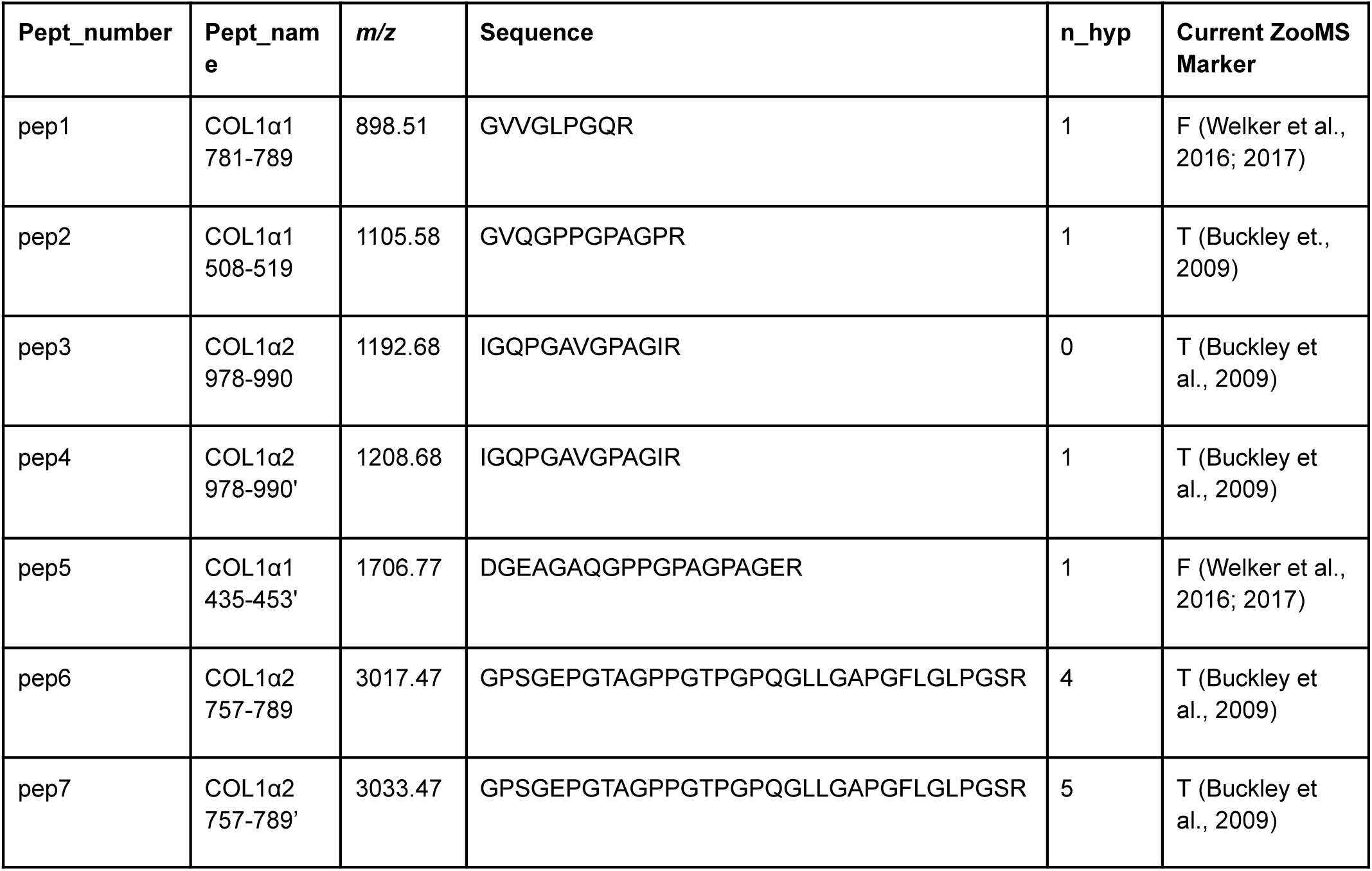
List of common peptides chosen to calculate MDS for the 9 taxonomic groups and their corresponding pept_number. Peptides are ordered by increasing *m/z*. References refer to the first publication of this peptide in ZooMS literature.

### Spectra preprocessing

The aim of the preprocessing is to extract the isotopic distribution of the selected peptides in order to calculate the extent of deamidation for each. This follows the same steps undertaken by the previous PQI method established by Nair and colleagues (2023). Our new MDS leverages the infrastructure provided by the Spectra R package (v4.4.0, (Rainer et al., 2022) to load and process the spectra. This allows for batch loading and processing of the spectra, overcoming the need for excessive memory, and allowing faster parallel processing, which presented a challenge in PQI. Once loaded, the spectra are smoothed using the Savitzky-Golay filter (Savitzky and Golay, 1964) using a half-window size of 6. Following this, the baseline is estimated using the Statistics-sensitive Non-linear Iterative Peak-clipping algorithm (SNIP) with 50 decreasing iterations (Ryan et al., 1988) on the weighted-moving-average smoothed (with half-window size 4) spectra to avoid the peak boundary artifacts introduced by the Savitzky-Golay filter (Schmid et al., 2022). This baseline is then subtracted from the Savitzky-Golay filtered spectra, which crucially is better at preserving peak shape. Peak picking is applied with a half-window size of 15 to detect local maxima. Noise is estimated using Friedman’s SuperSmoother (Friedman, 1984) and peaks above a signal-to-noise ratio (S/N) above 2 are retained. Pseudo isotopic distributions are extracted for each of the peptides by locating the monoisotopic peak plus 4 consecutive peaks at distance of 1 Da with a fixed tolerance of 0.4 Da and 50 ppm. Incomplete isotopic patterns with gaps or less than 4 peaks are discarded as in Nair et al. (2023). Finally, the intensities of each peptide’s isotopic envelope are normalized to its highest peak.

### Taxon specific peptide selection

Based on the number of isotopic peaks extracted for each selected peptide, users should further select peptides that either (i) have a high proportion of spectra with at least four isotopic peaks in the envelope, or (ii) show consistently low isotopic peak detection but are commonly observed across the selected species. In either case, users are expected to justify their peptide selection based on the detection profile and the biological or taxonomic relevance of the peptides.

### Estimation of peptide deamidation

We fitted the theoretical isotopic distribution of the non-deamidated and deamidated version of the peptides to the data using WLS regression and obtained the coefficients, γ_0_ and γ_1_, which represent the amount of each deamidation state respectively, calculated as:

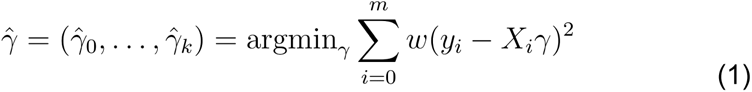

Where *k* is the number of glutamines, or deamidation states, in the peptide, and the number of isotopic peaks used. *y_i_* is the normalised intensity of the *i-th* peak in the observed envelope, *X_i_* is a row vector with the theoretical peaks values for the *i-th* position and γ a column with the coefficients. For the case of *k=1* from the estimated 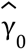 and 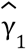, the fraction of deamidation is then

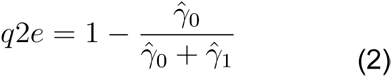

The weights, *w*, used in equation 1 are the signal-to-noise ratio and the inverse of the mass error of each peak of the isotopic series. The difference with the approach by Nair and colleagues (2023) is that here, the observed intensities (*y_i_*) are normalised for each isotopic envelope and therefore the scaling factor is not required.

### Quality control

Throughout the process of MDS we applied several quality control filters based on different measurements. First we removed the peptide envelopes from any sample in which an isotopic peak had a mass error above 0.4 Da. For WLS-q2e estimation, we calculate the reliability as the squared sum of the residuals and remove any peptide estimate from any sample with a reliability above 0.2. Finally, any WLS-q2e estimate below 0 or above 1 is discarded.

### MDS estimation

MDS is calculated as described in Nair et al. (2023). Briefly, a linear mixed effect model (LME) is fitted using the log*(q2e)* values as the response variable, the peptide as the fixed effect, and the sample and replicate as the random effects. The squared sum of errors of the previous WLS step (reliability) is used as a weight or scaling factor for the residual variances (see Nair et al., 2023). The MDS is the sample-level random effect of the model, which is extracted after the model is fitted. In sum, the whole workflow of MDS is illustrated by **Figure 1**.

**Figure 1.**
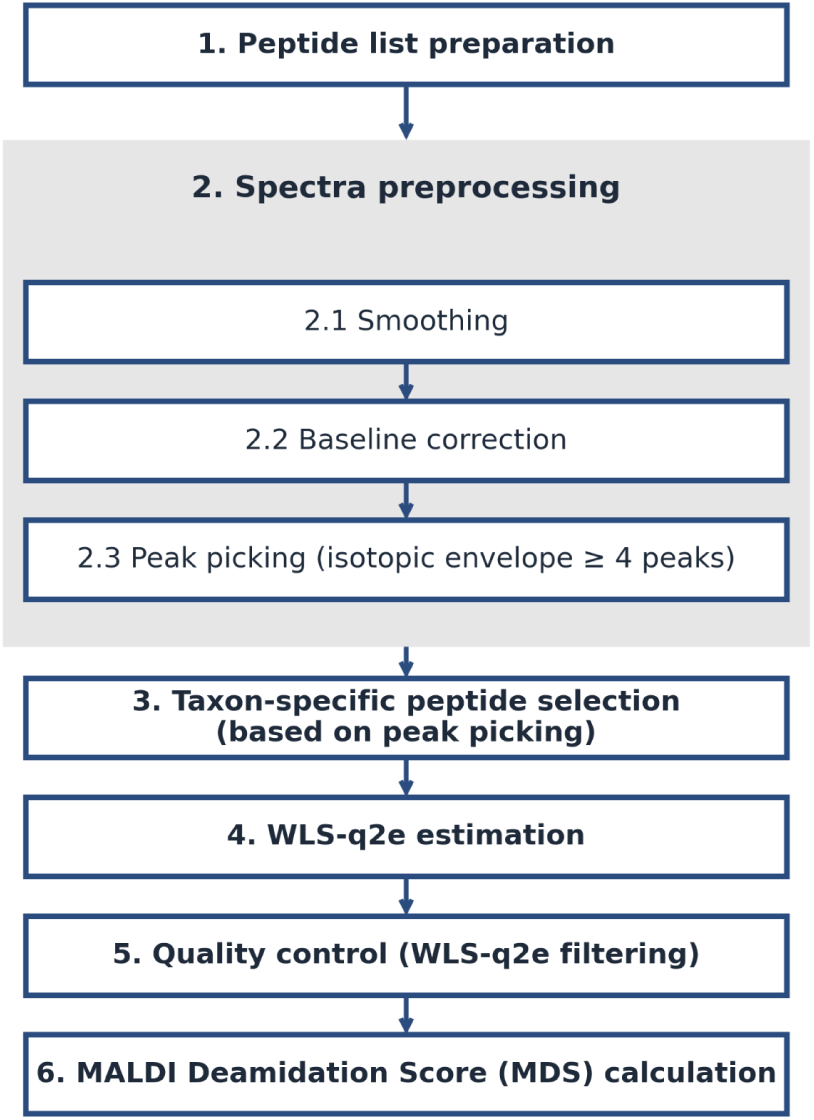
Workflow for calculating MDS. The process begins with peptide list preparation, followed by spectra preprocessing including smoothing, baseline correction, and peak picking (requiring isotopic envelopes with ≥4 peaks). Based on the peak picking results, taxon-specific peptides are selected. WLS q2e estimation is then performed, followed by quality control (QC), and finally, the MDS is calculated.

### Cluster analysis as another line of evidence

We performed a hierarchical cluster analysis of the *Bison/Yak* samples. For this, samples were independently preprocessed. Samples were first smoothed using a Savitzky-Golay filter with half-windows size 8, then baseline was estimated using the SNIP algorithm with 50 decreasing iterations. Local maxima were detected using a half-window size of 15. Peak centroid *m/z* position was corrected using the weighted average of the 10 adjacent decreasing points. Finally, peaks above SNR 3 are kept. Monoisotopic peaks were detected using the algorithm provided in ‘MALDIquant’ (Gibb, 2024; Gibb and Strimmer, 2016, 2012), based on predicted theoretical isotopic envelopes (Breen et al., 2000). We used a minimum correlation of 0.1 between observed and theoretical envelopes, so deamidated peptides are included, and 100ppm matching tolerance. Furthermore, peak spectra were aligned using an internal reference of peaks appearing in at least 20% of the *Bison/Yak* samples, following the algorithm proposed in MALDIquant (Gibb and Strimmer, 2016). Finally, peaks *m/z* values closer than 0.1 are binned, and replicates merged. Finally, the peak spectra are arranged in a feature matrix, with samples in the rows, *m/z* values in the columns and filled with intensity values.

The contaminant list provided by SpecieScan (Végh and Douka, 2024) (https://github.com/mesve/SpecieScan/blob/main/Reference_databases/contaminants_list.csv) was used to filter out possible contaminants. Differential presence of contaminants can be a source of difference between batches.

The matrix features were standardised to mean one and variance zero. A hierarchical clustering was carried out using euclidean distance and the ‘ward.D2’ agglomerative method.

## Results

### Improvement of MDS

#### More efficient preprocessing

We tested the processing times on the *Bison/Yak* samples (2178 spectral files according to the metadata from Brown et al., 2021), and found that GA-q2e, without considering any manual filtering or replicate averaging, was about 3 and 22 times slower than the complete PQI and MDS pipelines, respectively (**Table 2**).

**Table 2:**
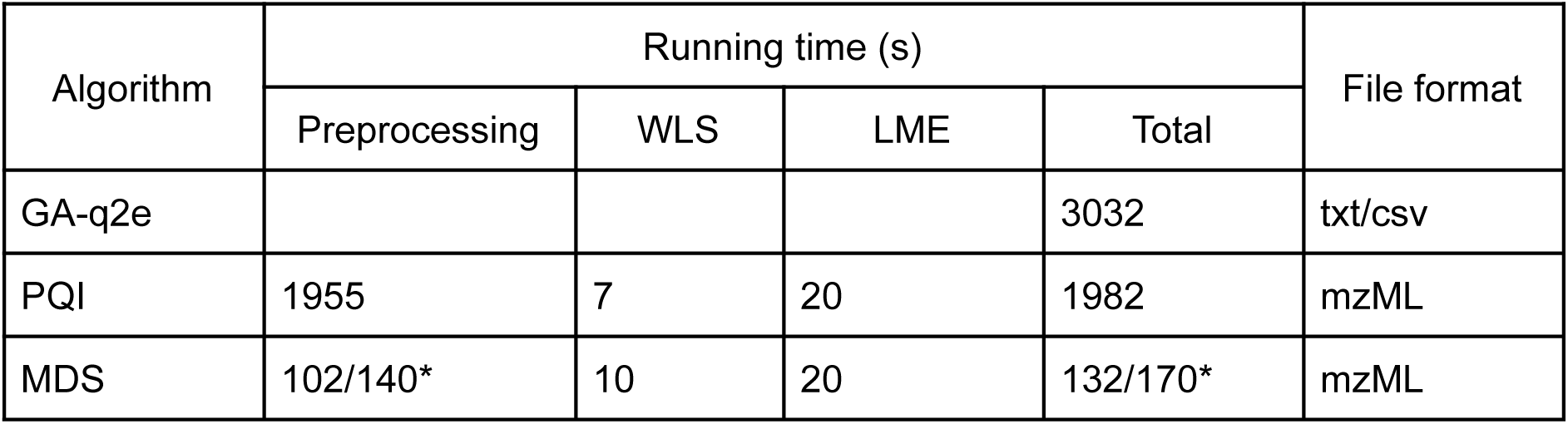
Comparison of running times for MDS, GA-q2e, and PQI on the Bison/Yak dataset. The total processing time for GA-q2e was 3032 seconds, excluding the time required for converting mzML files to TXT or CSV formats. For MDS, the reported preprocessing time includes all spectrum and aggregate files from the triplicate measurements (*).

The GA-q2e method relies on parsing text-based CSV files, a computationally demanding process compared to reading spectral data stored in binary format (mzML). Additionally, GA-q2e employs slower preprocessing algorithms for signal-to-noise (S/N) estimation and peak detection. Most significantly, the genetic algorithm underlying GA-q2e greatly increases computation time compared to the WLS method, wherein values can be calculated directly using an analytical expression. In turn, MDS achieves substantial improvements in computational efficiency relative to the earlier PQI method by leveraging the Spectra package infrastructure (Rainer et al., 2022) for handling MALDI-TOF spectral data. Thus, the most notable efficiency gains among these methods occur in the preprocessing step of MDS.

#### Normalization of isotopic series

The new MDS tool formats the extracted peptide isotopic peaks data to use standard R model functions (like ‘lm’) for peptide WLS-q2e estimation step, instead of an ad-hoc WLS fitting function, as it was implemented on the MALDIpqi package (for the PQI method). This allows more flexibility and potentially the use of different models or fitting functions within the R environment. Furthermore, the observed intensities within each isotopic envelope are internally normalised to the same scale as the theoretical isotopic envelopes, which removes the need for an extra scaling factor as was necessary for the WLS step in the PQI method.

### Differential detection of peptide isotopic envelopes

The estimation of q2e values by WLS in the MDS pipeline relies on the accurate detection of the selected peptide isotopic envelopes. From the peak picking step of our preprocessing pipeline, we observed differential detection of peptides, including the same or equivalent peptide marker across taxa (**Figure 2**).

**Figure 2.**
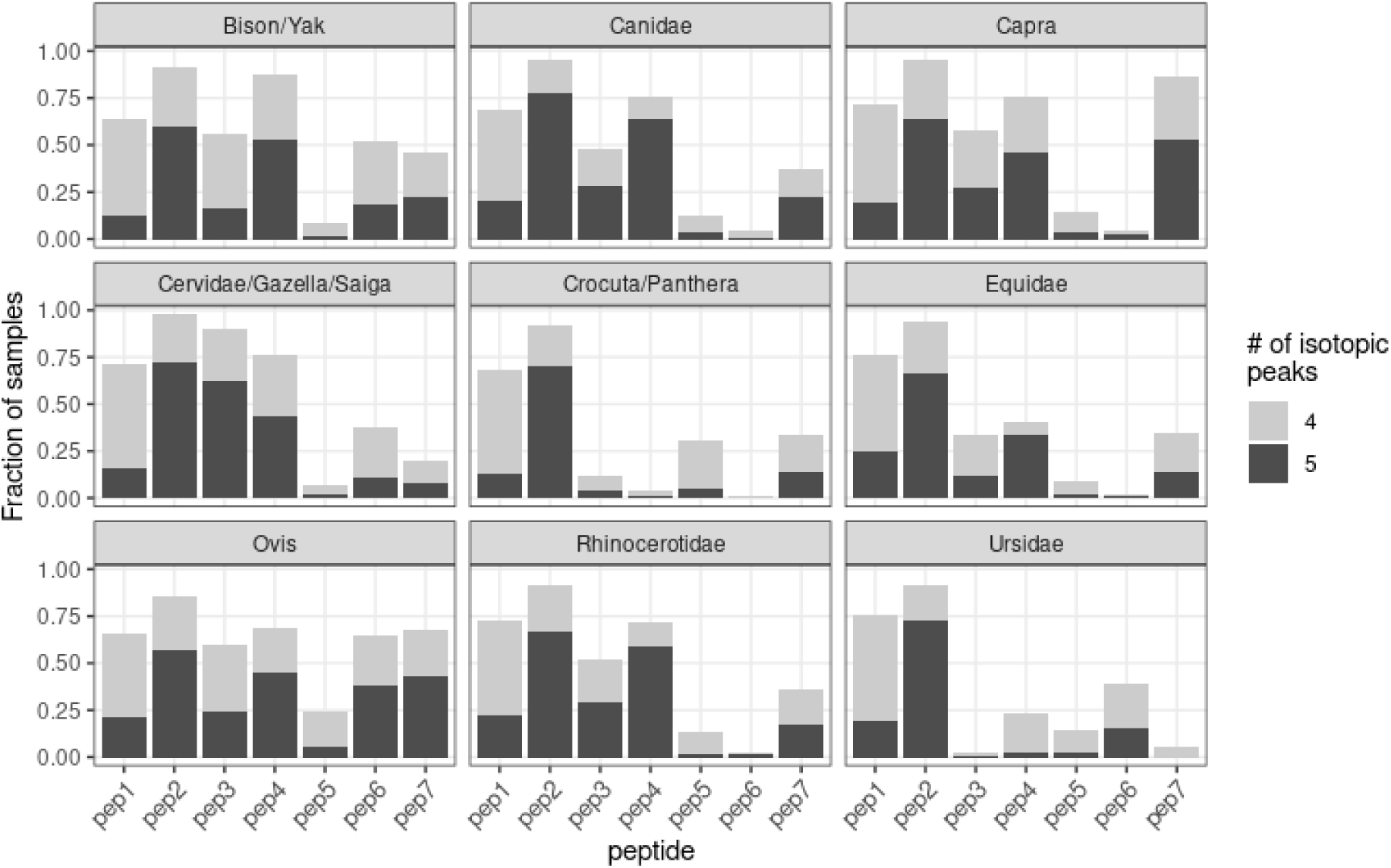
Fraction of samples in each taxon where each peptide was detected with four or five isotopic peaks. Peptides with at least four isotopic peaks are eligible for inclusion in MDS-based deamidation estimation. Color shading indicates the number of peaks detected (light gray = 4, dark gray = 5).

Pep1 (COL1α1 781-789) and pep2 (COL1α1 508-519) are reliably detected with isotopic envelopes of at least four peaks across most taxa. Pep3 and pep4 correspond to the COL1α2 978-990 marker and its fully hydroxylated version. These peptides are consistently recovered across all herbivores except *Equidae*. Among carnivores, pep3 and pep4 are only reliably observed in *Canidae*, while the homologous sequence in *Crocuta/Panthera* (TGHPGSVGPAGVR) lacks a glutamine residue and is therefore excluded from MDS analysis. Even so, only about 25% of *Crocuta/Panthera* spectra exhibit a detectable isotopic envelope for either the hydroxylated or non-hydroxylated variant, indicating its low detection rates in MALDI-TOF MS. A similar pattern is observed in *Ursidae*, where the pep3 and pep4 markers were excluded due to the absence of a glutamine. Furthermore, when detected, this peptide in *Ursidae* predominantly appears in its hydroxylated form.

Pep6 and pep7 correspond to COL1α2 757-789 and its hydroxylated variant. In *Bison/Yak*, *Ovis*, and *Cervidae/Gazella/Saiga*, pep6 overlaps with a common conserved collagen peptide for bovid (GPPGASGAPGPQGFQGPPGEPGEPGQTGPAGAR; COL1α2 89–130) (Janzen et al., 2021) and is frequently detected. However, for all other taxa in which pep6 does not overlap with COL1α2 89–130 (except *Ursidae*), typically only the hydroxylated version is detected, and the non-hydroxylated version is rarely observed. This suggests that pep6 sequences identified for *Bison/Yak*, *Ovis*, and *Cervidae/Gazella/Saiga* likely originate predominantly from the non-marker peptide. *Ursidae* exhibits a reversed pattern for pep6 and pep7, likely because the corresponding peptide sequence for *Ursidae* (GPSGEPGTAGPPGTAGPQGIIGAPGIIGIPGSR) contains only four prolines in the Y-position that can be easily hydroxylated (which generally tend to be fully hydroxylated), whereas in other taxa, the equivalent peptide contains five hydroxyproline residues.

Pep5 (COL1α1 435-453) is common across all taxa but is poorly represented in the Denisovan dataset. For instance, only around 25% of the pep5 detected for *Ovis* and *Crocuta/Panthera* contain a clear isotopic envelope with at least four peaks; notably, this detection rate is already the highest among all taxa. Nevertheless, despite its relatively low spectral representation, pep5 was included in the MDS analysis because of its taxonomic ubiquity and its prior application in q2e-based deamidation estimations (Welker et al., 2017).

Therefore, based on the peak picking results, peptides were further refined from the preselected peptide list (**Table 1**) for downstream MDS estimation. Specifically, the selected peptides included pep1, pep2, pep3, pep4, pep5, and pep7 for *Bison/Yak*, *Ovis*, *Capra*, *Cervidae/Gazella/Saiga*, *Rhinocerotidae*, *Equidae*, and *Canidae*; pep1, pep2, pep5, and pep7 for *Crocuta/Panthera*; and pep1, pep2, pep5, and pep6 for *Ursidae*.

### WLS-q2e peptide filtering

Following our quality control procedure, which filters out unreliable peak data based on WLS sum of squared residuals, WLS-q2e values were generated for each of the samples using MDS based on the selected peptides. Samples with at least two valid WLS-q2e values were retained for further analysis. This filtering resulted in 2,169 samples being included in the final MDS dataset, comprising 718 *Bison/Yak*, 750 *Cervidae/Gazella/Saiga*, 306 *Equidae*, 164 *Rhinocerotidae*, 63 *Canidae*, 63 *Capra*, 52 *Ursidae*, 27 *Ovis*, and 26 *Crocuta/Panthera*.

Pep1 and pep2 are the two most reliable peptides. For pep2 specifically, WLS-q2e values were generated in 2,137 samples, of which 2,109 passed quality control and were retained for downstream analysis; slightly fewer pep1 values passed the quality control. Peptides with higher *m/z* tend to lose more values during the filtering. Interestingly, among the marker pair pep3 and pep4, pep3 showed the most significant data loss (>50%) for *Cervidae/Gazella/Saiga* and *Canidae*, whereas pep4 exhibited greater data loss for *Equidae* and *Rhinocerotidae*. **Figure 3** summarizes the number of WLS-q2e values generated and retained for each peptide across taxa.

**Figure 3.**
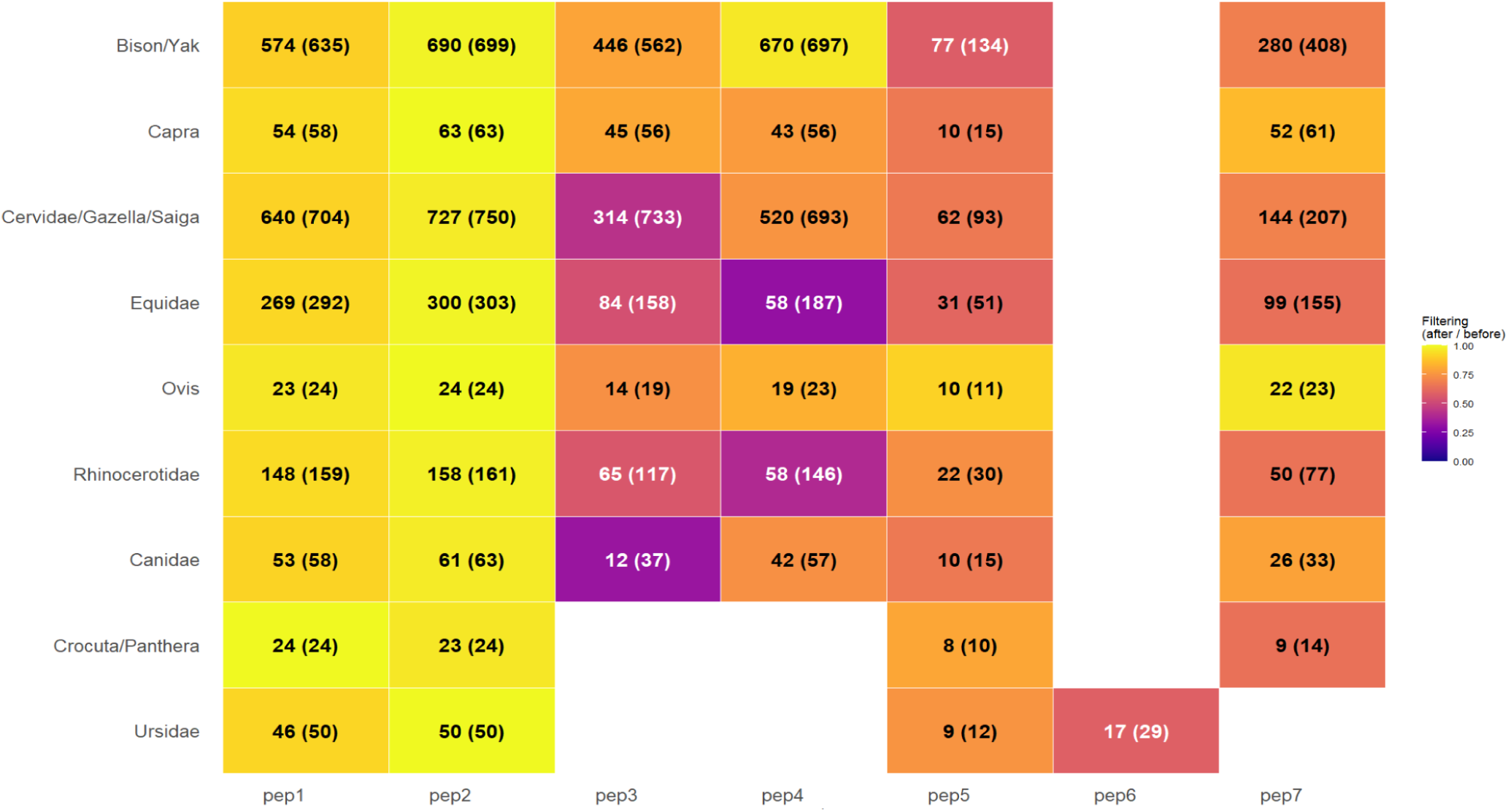
Peptide counts per taxon before and after quality filtering. Each cell indicates the number of WLS-q2e estimates retained for each peptide. Values in parentheses represent the total counts before filtering.

### Peptide level deamidation

To evaluate the performance of the WLS regression on the selected peptides, we compared WLS-q2e values from all analyzed peptides against GA-q2e values derived from pep2. Overall, a strong positive correlation was observed across taxa, demonstrating broad consistency between the two approaches (**Figure S1**). However, we still observe substantial variability among taxa and peptides (**Figure 4**). Pep1 and Pep2 are the most frequently recovered peptides in the MDS analysis. Across all taxa, Pep1 consistently shows higher deamidation levels than Pep2, with median values generally exceeding 0.5, whereas the median for Pep2 for cases is below 0.5 (**Figure 4**). This suggests that within the analysed dataset, Pep1 is deamidated more rapidly than Pep2.

**Figure 4.**
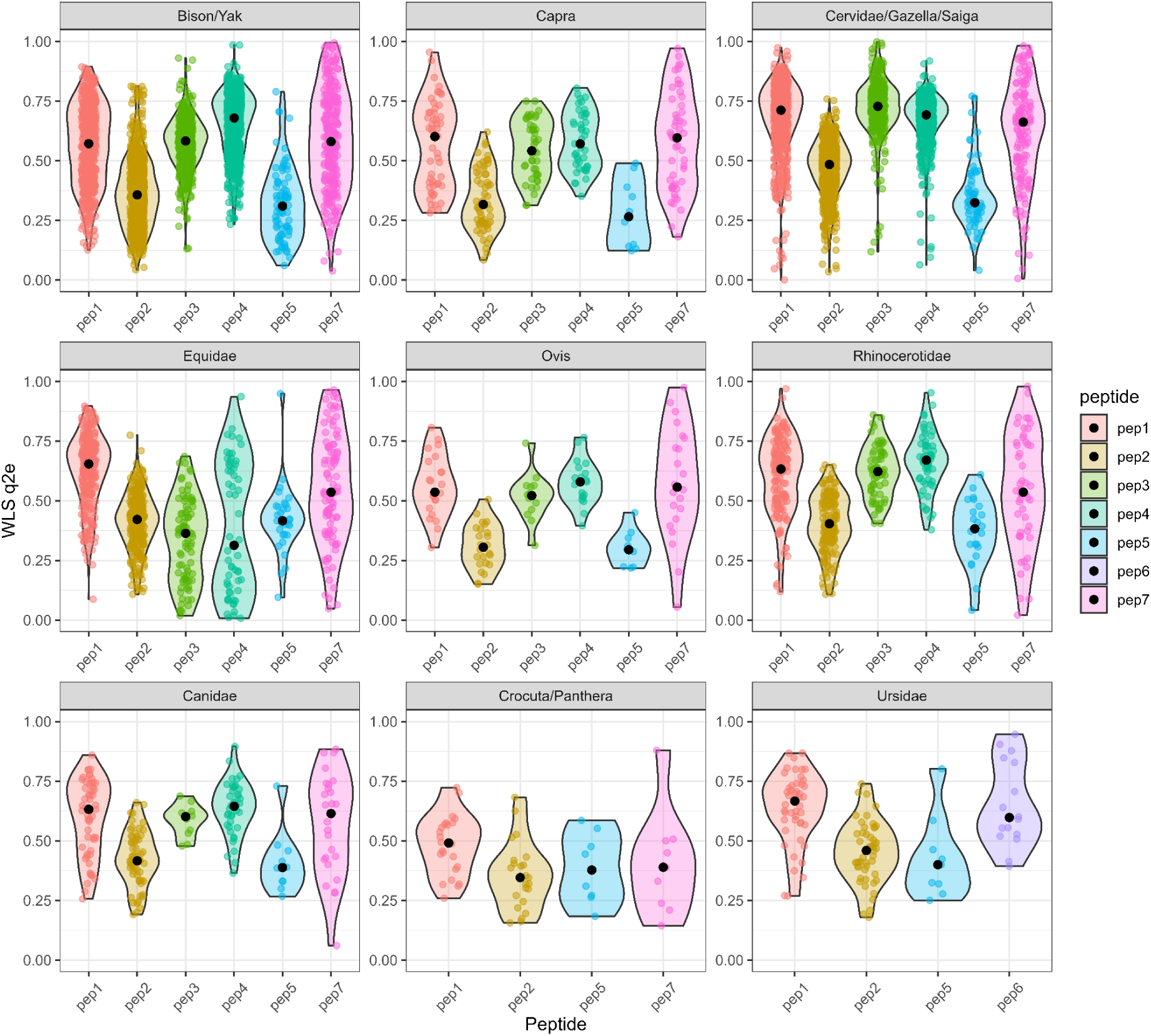
Violin plots showing the distribution of WLS-q2e values across individual peptides (pep1–pep8) for all the nine taxa. Each panel represents a taxonomic group, with individual peptide distributions coloured consistently across panels. Median values are shown by the black dots.

The ZooMS marker pair pep3 and pep4 also show elevated deamidation values compared to pep2, with median always higher than 0.5 but with subtle variation between the two. In most taxa, pep4 is more deamidated than pep3 in terms of the median, except in *Cervidae/Gazella/Saiga*, where the reverse pattern is observed. In *Equidae*, pep4 displays a bimodal distribution, with one group of data clustered below 0.5 and one group above, potentially indicating the presence of two distinct sample preservation conditions. This will be discussed further below.

Pep5, although detected in fewer samples, shows relatively low deamidation levels (mostly <0.5) across all taxa, suggesting a slower deamidation rate for this peptide similar to pep2. In *Ursidae*, pep6 stands out as the most deamidated peptide, while pep7, present in multiple taxa, spans nearly the entire range (0–1), indicating substantial variation in degradation levels and possibly reflecting peptide-specific detectability on the MALDI spectrum.

It is clear that pep1 and pep2 are the two best candidates among the selected peptides for the deamidation estimation due to their consistent presence and their well-defined isotopic envelopes in MALDI-TOF spectra. Among these two, pep2 is better when dealing with assemblage covering deep time as it deamidates the slowest. However, we do not know if this is specific to the assemblage we are testing or a universal phenomenon. In addition, other factors such as the physicochemical properties of the peptides may affect their efficiency in extraction and detection during the lab process. For example, pep1 and pep2 may more easily bond with the C18 resin used in peptide purification in standard ZooMS protocols. They may also co-crystallise particularly efficiently with the matrix solution used to facilitate ionisation of the peptides to enable detectability in the TOF mass analyzer (Körsgen et al., 2016). The large variation for heavier peptides such as pep6 and pep7 may be the product of overlapping irrelevant peptide peaks on the spectra.

### Diachronic Patterns in Peptide Deamidation

Across stratigraphic layers (**Figure 5**, **Figure S2**), most peptides exhibit a general U-shaped pattern in WLS-q2e values, with decreasing deamidation from the oldest layers (e.g., layer 15 and, where available, 17.1) toward mid-sequence layers, followed by an increase in the youngest layers (9.3 to 9.2). However, both inter-peptide and inter-taxon variability are evident. Pep1 and pep2, which have the highest number of observations, clearly follow this U-shaped trend across most taxa. Pep3 and pep4 display similar distributions across layers when given sufficient sample size Pep5, pep6, and pep7 lack sufficient data to establish robust trends across all taxa. Nevertheless, for more abundant taxa such as *Bison/Yak*, pep5 and pep7 show similar trends compared to the well detected pep1 and pep2. However, the high variation of pep7 values, including many near-complete deamidation values close to 1, makes subtle interlayer differences difficult to discern.

**Figure 5.**
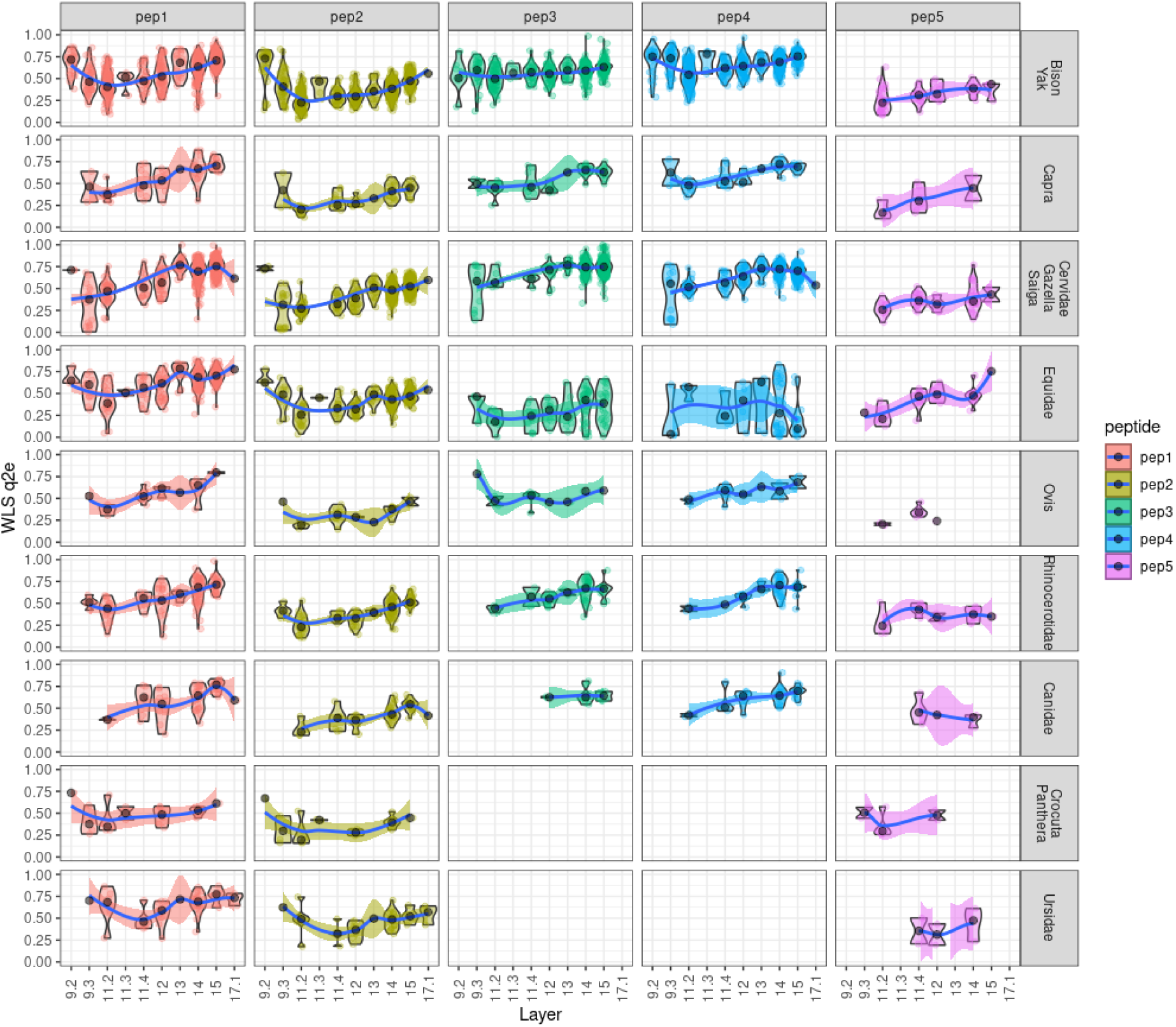
Peptide deamidation patterns across layer and taxa on pep1 to pep5. For the same plot on pep6 and pep7, see supplementary **Figure S2**.

Interestingly, peptide-specific deviations are observed within certain layers. For example, in *Bison/Yak*, the behavior of pep1 and pep2 diverges in layer 11.3 and layer 13 (**Figure 6**). For layer 11.3, although the sample size is small (*n*=5), all five samples show WLS-q2e values for pep2 that are higher than the median values of both adjacent layers. This pattern is not observed in pep1, for which the WLS-q2e values instead show a gradual decreasing sequence from layer 11.4 to layer 11.2. Similarly, pep3 and pep4 also exhibit locally elevated values in layer 11.3, although based on even fewer observations (**Figure 5**). On the contrary, pep2 shows a gradual decreasing trend from layer 14 to layer 11.2 with no layers show statistically significantly higher values in between, while pep1 shows elevated WLS-q2e values in layer 13 which are higher than the values in layer 12 and layer 14, although not significantly for the latter. The elevated WLS-q2e values for pep1 in layer 13 are also associated with a lower sample size compared to pep2 (*n*=18 vs. *n*=22), though the same association is not seen in layer 11.3. A similar, though subtler, pattern is evident in *Cervidae/Gazella/Saiga*. While no samples were recovered from layer 11.3, pep1 exhibits significantly elevated WLS-q2e values in layer 13 compared to layer 12, and slightly elevated values compared to layer 14, although the latter difference is not statistically significant. In contrast, pep2 in layer 13 shows only a slight but statistically significant elevation compared to layer 12 and is similar to layer 14 by visual assessment (**Figure 6**).

**Figure 6.**
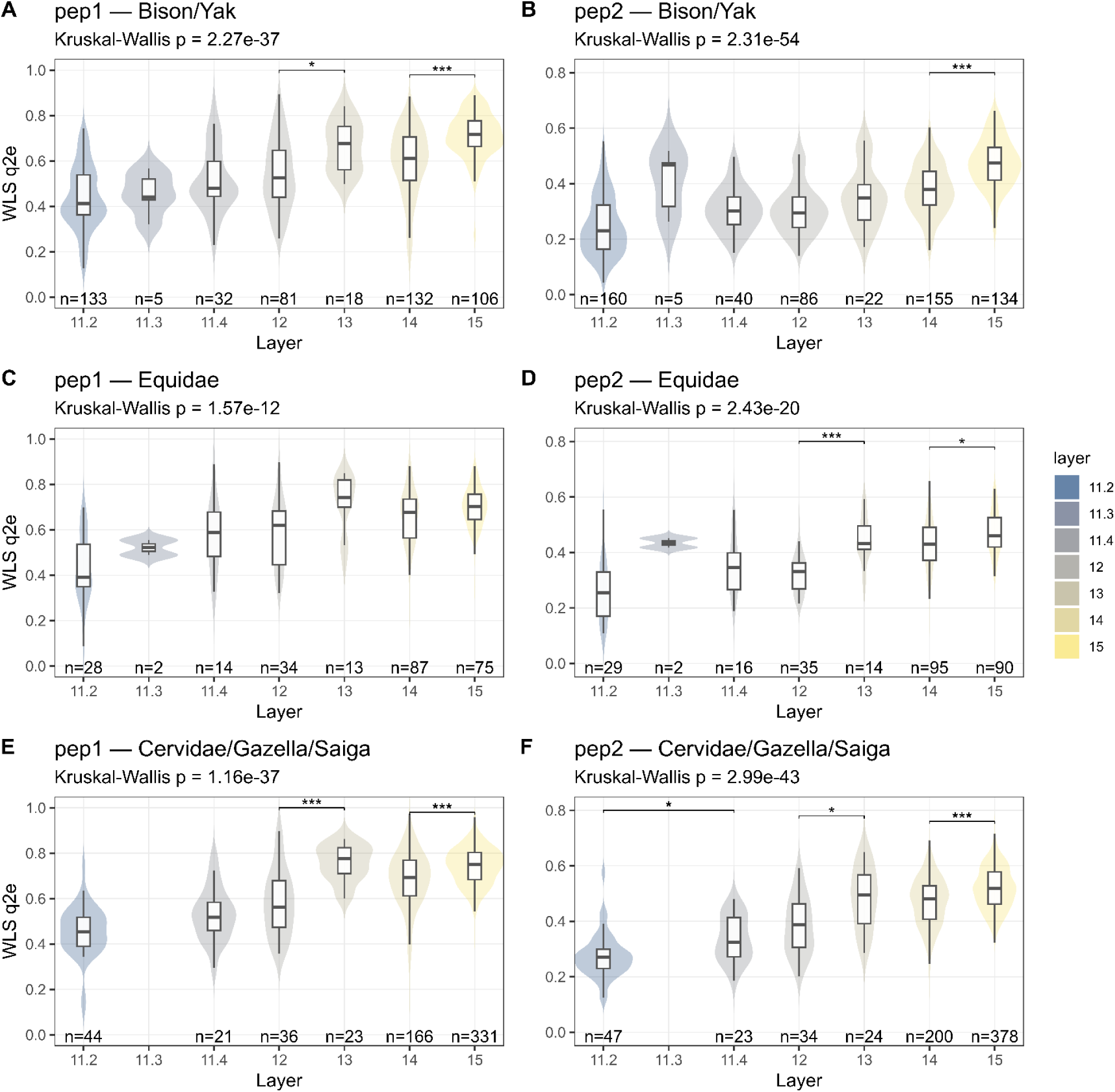
Distributions of WLS q2e for pep1 and pep2 across layer 15-11.2. Panels are arranged by species (rows) and peptides (columns): A–B, *Bison/Yak*; C–D, *Equidae*; E–F, *Cervidae/Gazella/Saiga*; left column = pep1, right column = pep2. For each layer (11.2–15), violin plots show the kernel density, with overlaid boxplots indicating the median and interquartile range (whiskers = 1.5*IQR); individual points are not displayed. Per-layer sample sizes (*n*) are printed near the x-axis. Layers with no observations for a given species (e.g., 11.3 in some groups) remain on the x-axis but have no violin/box. Panel subtitles report the global Kruskal–Wallis *p*-value across layers for that species–peptide combination. Pairwise tests are limited to adjacent layers (Wilcoxon rank-sum with Bonferroni correction): *p* < 0.05 (\**), p < 0.01 (**), p < 0.001 (\*\*\**); non-significant contrasts are omitted.

*Equidae* appears to differ from this trend associated with layer 13. For pep2, the values in layer 13 are significantly higher than those in layer 12, a pattern not observed for pep1. However, this discrepancy might be attributed to the relatively small sample size available for *Equidae* in layer 13. In layer 11.3, only two samples were recovered for both peptides; notably, the pep2 values of these two samples exceed the 75th percentile of values from the two neighboring layers, whereas both pep1 values fall well below the median of layer 11.4, mirroring the pattern observed in *Bison/Yak*. (**Figure 6**) In other taxa, limited sample sizes constrain a clear interpretation on the deamidation trends of each peptide, but general U-shape trends across all layers remain broadly consistent (**Figure 5**, **>Figure S2**).

#### Equidae

As noted in **Figure 4**, we observed an unusual distribution of WLS-q2e values for *Equidae*, particularly for pep4. Upon closer inspection, this anomaly appeared to be driven by a clear bimodal distribution, separating the samples into two distinct value groups. To assess whether this pattern was caused by batch effects during spectrum acquisition, we plotted WLS-q2e values of pep4 across different batches (**Figure 7A**). The grouping persisted across batches, suggesting that batch-specific artifacts were not the primary cause.

**Figure 7.**
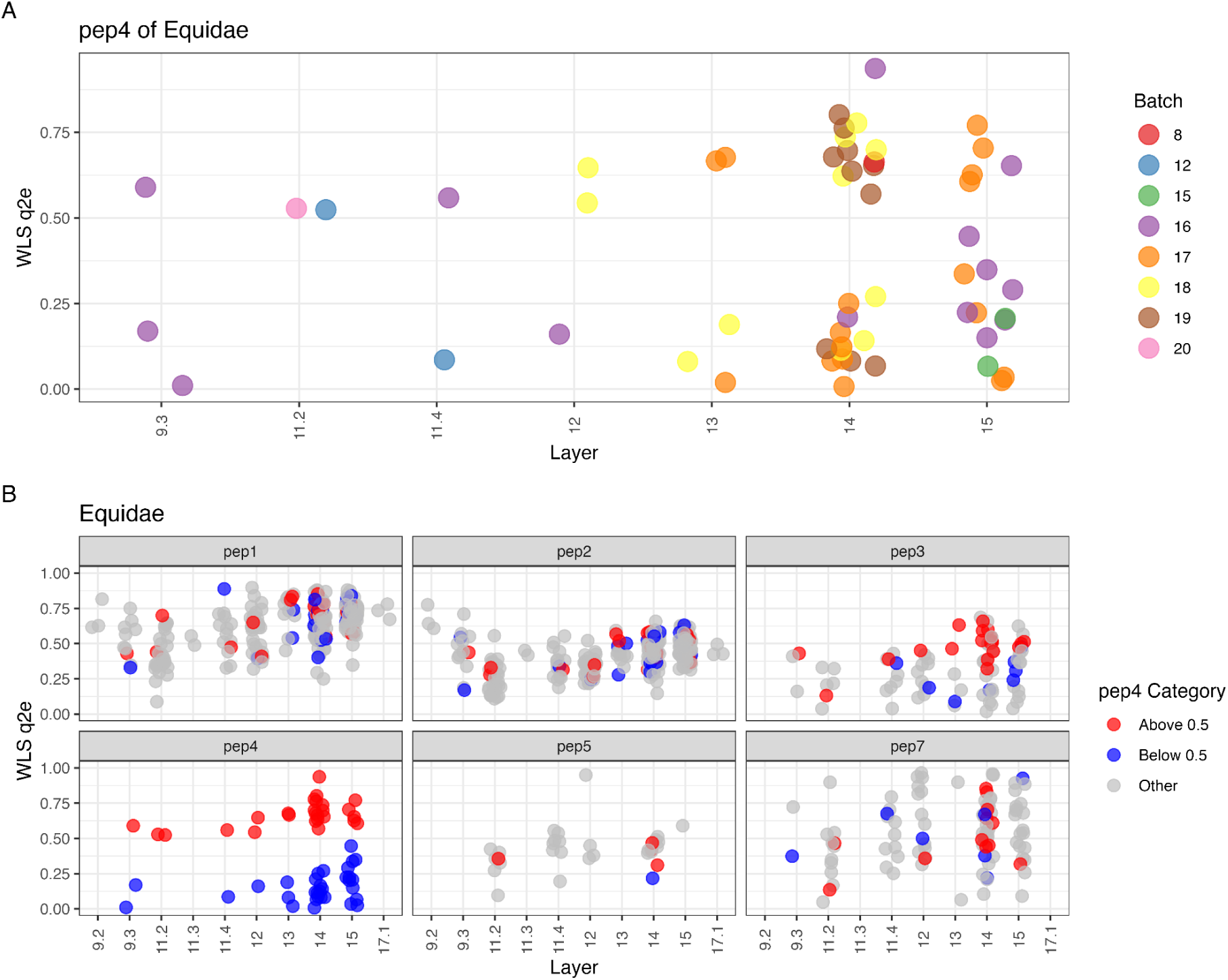
**A)** Distribution of WLS-q2e values for pep4 in *Equidae* across batches, **B)** WLS-q2e values of other peptides in *Equidae* samples categorized by pep4 WLS-q2e above or below 0.5 group.

To further investigate this phenomenon, we mapped the two pep4-defined groups (above vs. below WLS-q2e = 0.5) onto the WLS-q2e distributions of other peptides from the same samples (**Figure 7B**). This revealed no corresponding groupings in most peptides, except pep3, where a weak co-pattern was observed but the limited sample size precludes firm conclusions on pep3.

Using the spectral inspection feature integrated into the MDS preprocessing workflow, we retrieved the original spectra of samples exhibiting a grouping effect on pep4. The spectral examination revealed that the group with lower WLS-q2e on pep4 had an overlapping peptide isotopic envelope that shifted its detected envelope towards having a lower deamidation value (**Figure 8**). This phenomenon worryingly passed the quality control based on the sum of the squared errors of the residuals of the WLS fit. In principle, this should identify and remove distorted isotopic envelopes with overlapping peaks. In this case, using WLS regression models would not produce accurate q2e values and therefore have large errors. Why some *Equidae* samples have this overlap is unknown to us, as it appears to occur across batches (**FIgure 7A**).

**Figure 8.**
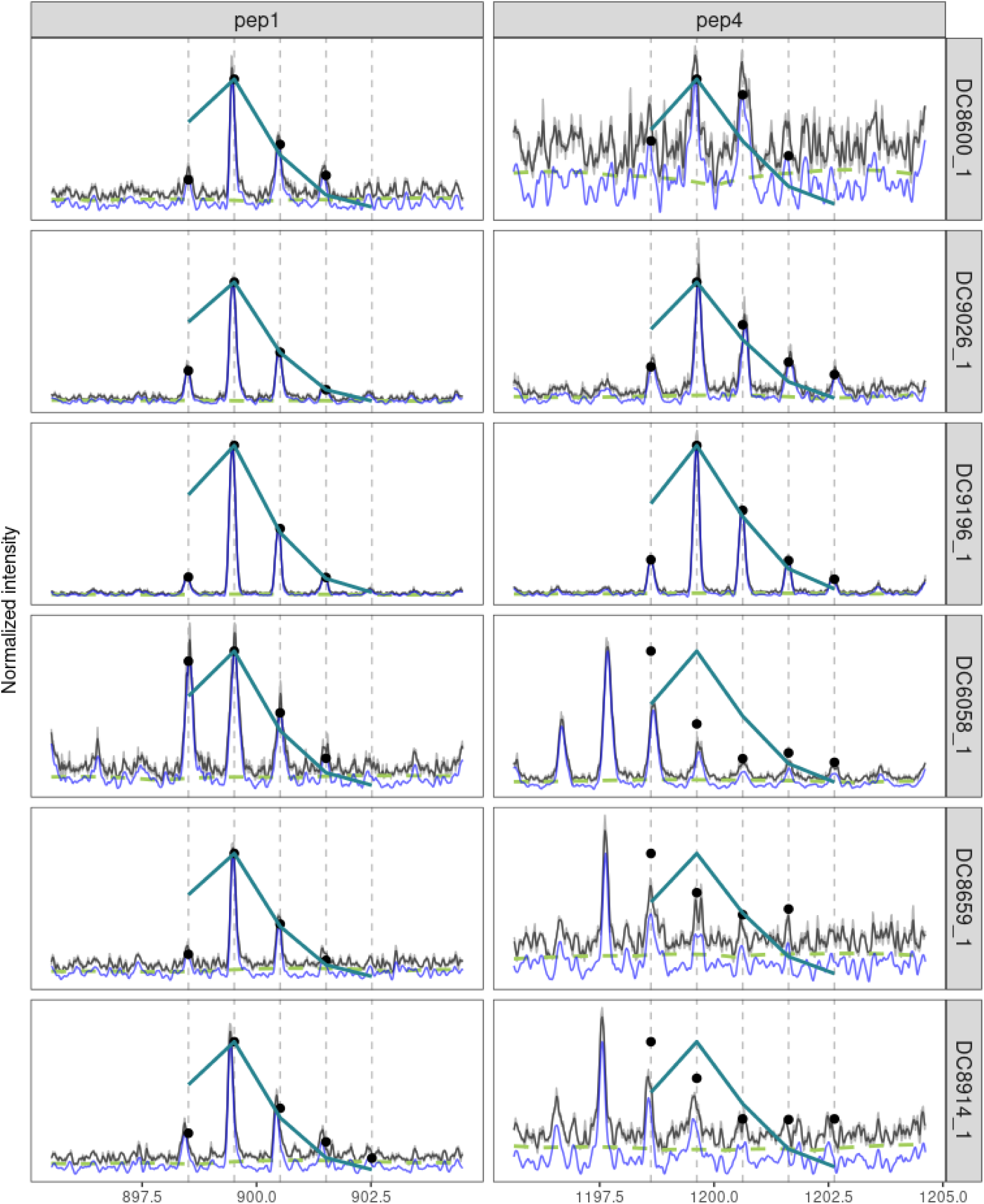
Pep 1 and 4 of six different *Equidae* spectra. The top three are spectra with WLS-q2e above 0.5 and the bottom three below 0.5. The grey lines are the raw and smoothed spectra and the blue line the baseline corrected. The black dots are the normalised isotopic peaks and the turquoise line a theoretical shape of a peptide with q2e 0.5.

### MDS and GA-q2e

The overall MDS of each sample was calculated by integrating peptide-specific WLS-q2e values using a linear mixed-effects model (LME). Diachronic trends of MDS (**Figure 9**) closely mirror those of GA-q2e from Brown et al., 2021 (**>Figure S3**). Both methods show a decline in deamidation from Layer 15 to Layer 11.4, with a minor rise in Layer 13, followed by another decline at Layer 12. A small local increase is observed in Layer 11.3, though based on limited data. Both MDS and GA-q2e reach their minimum in Layer 11.2, after which deamidation levels rise sharply from layer 9.3 to 9.2 (**Figure 9**). This similarity is further supported by a strong linear correlation between MDS and GA-q2e values across samples (**Figure 10**). A global linear regression yields a slope of 1.05 (*R²* = 0.62, *p* < 0.0001), indicating a strong and statistically significant agreement between the two methods, albeit with a slightly elevated intercept, such that MDS values start at a higher level.

**Figure 9.**
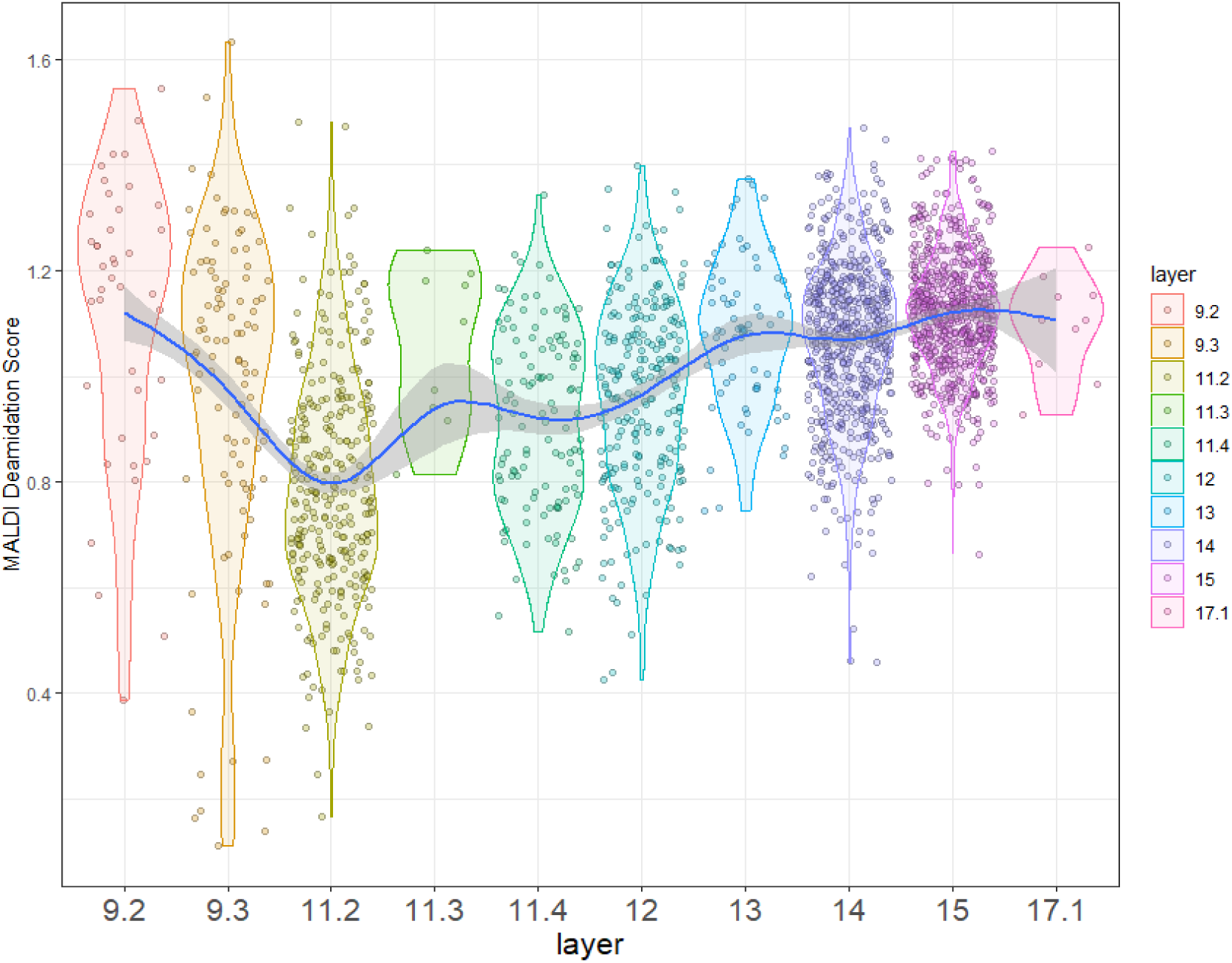
Diachronic patterns of MDS and GA-q2e across stratigraphic layers.

**Figure 10.**
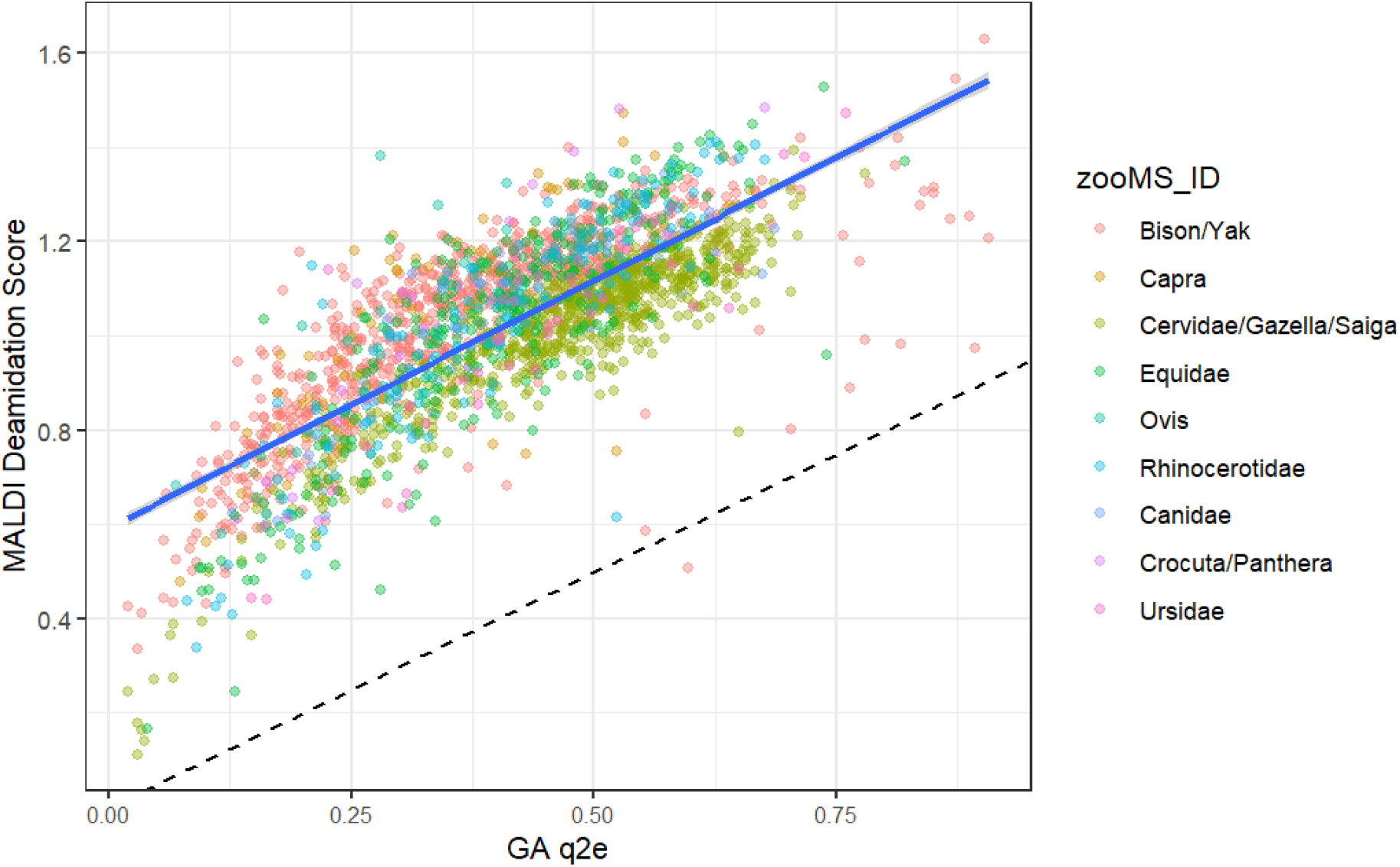
Correlation between MDS and GA-q2e across taxa.

**Figure 11.**
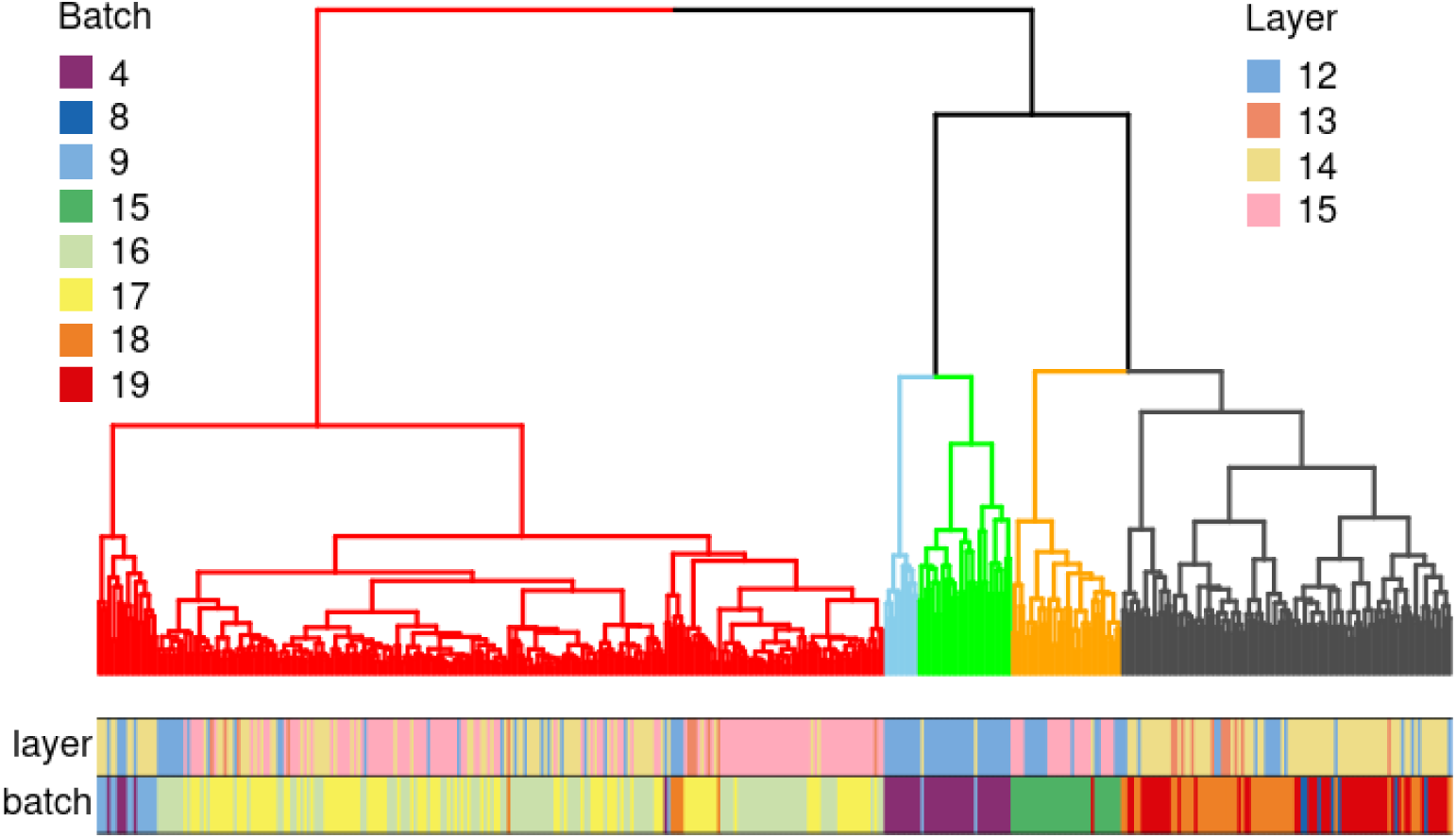
Hierarchical clustering of the Bison/Yak samples and layers 15 to 12.We show the five most distinct clusters, with branches colored according to them. Each sample’s layer and batch are indicated in the two bottom colorbars.

### Clustering analysis on whole spectra of *Bison/Yak*

We conducted hierarchical cluster analysis on all spectra of *Bison/Yak* used in this study to explore whether other modifications, particularly hydrolysis, might have influenced the spectral patterns. Samples from layers 15 to 12 were selected because these layers exhibit a gradual decrease in deamidation levels (**Figure 5**, **Figure 9**). If intensive hydrolysis had occurred, we would expect to see higher peaks corresponding to shorter peptides and lower peaks for longer peptides, as longer chains would have been broken down during deposition. However, when plotting the spectra with associated metadata (layer, batch, and taxa), it became clear that clustering was overwhelmingly dominated by batch effects.

## Discussion

### Fast, Streamlined, and Flexible

Although the resolution of MALDI-TOF derived ZooMS spectra limits the precise quantification of post-translational modifications (Bray et al., 2023; Simpson et al., 2019), deamidation estimations produced by the GA-q2e method have been already applied for assessing relative chronology, contamination, and taphonomy. A niche for ZooMS studies is the analysis of large and indeterminate osseous assemblages either to complement zooarchaeological records (Lee-Michaelis et al., n.d.; Sinet-Mathiot et al., 2019); or as a pre-screening method of identifying hominin remains (Brown et al., 2016; Devièse et al., 2017; Mylopotamitaki et al., 2024; Welker et al., 2016) or the remains of rare fauna (Buckley and Herman, 2019; Hofman et al., 2018). These projects have generated a vast repository of ZooMS data at such a rapid speed that meta-analyses at site, regional, and time-series scales are now possible. This accumulation of ZooMS spectra will be further accelerated as laboratory robotics (Oldfield et al., 2025) and automated spectral post-processing advances (Gu and Buckley, 2018; Hickinbotham et al., 2020; Végh and Douka, 2024). Our benchmarking of processing times for MDS, GA-q2e, and PQI demonstrates that MDS significantly outperforms GA-q2e in terms of speed. It also surpasses PQI, which is its base model.

A significant portion of the processing speed improvement comes from the elimination of numerous time-consuming data preparation steps required before running the deamidation model, as well as from streamlining the overall workflow. These time savings are often masked and difficult to quantify directly. For example, sampling strategy coupled with GA-q2e analysis often involves manual filtering of samples with failed triplicates (Brown et al., 2021) or additional processing such as averaging spectral triplicates (Huan et al., 2024), making GA-q2e difficult to scale with the potential and growing need for meta-analysis and also loose valuable information from the differences within triplicates. Although PQI shares the same underlying statistical structure, it is less streamlined and requires additional time to execute each step of the workflow. MDS overcomes all these problems by integrating data preprocessing, quality control, and deamidation estimation into a single, automated pipeline. This streamlined design minimizes manual intervention and significantly improves overall processing efficiency, making it particularly well-suited for high-throughput analysis of large datasets.

MDS offer great flexibility for the users wishing to work on multiple taxa with different targeted peptides. The configuration of peptide lists is significantly simplified in MDS, allowing users to tailor taxa-specific peptide sets within a single Excel sheet that incorporates both *m/z* and sequence information. Although GA-q2e also permits user-defined peptide lists, its format includes only mass and sequence data without taxonomic specificity. This makes it difficult to process datasets involving multiple taxa with distinct peptide sets, unless further modifications are made to the underlying C code (Wilson et al., 2012). MDS also introduces rigorous preprocessing steps that enable more refined screening of usable peptides, in contrast to GA-q2e, which directly outputs deamidation values. We observed that peptides with higher molecular masses are more frequently filtered out, possibly due to lower spectral visibility or overlapping isotopic envelopes with non-target peptides. The observed difference in filtering rates between non-hydroxylated and hydroxylated peptide versions in certain taxa may be attributed to peptide overlapping effects in taxa-specific regions. With MDS’s built-in quality control features based on mass errors, SNR or reliability, it is now feasible to map peptide visibility more systematically and to provide informed guidance on the selection of suitable peptides for deamidation analysis using MALDI data and open doors for further analysis to inspect performance of individual peptides in MALDI-TOF MS.

### WLS-q2e reveal nuance pattern obscured by single peptide

A key feature of MDS is that it, for the first time, formalizes the output of WLS-q2e on each individual peptide selected per taxon. This enabled us to detect subtle fluctuations in deamidation rates between groups such as *Bison/Yak*, *Cervidae/Gazella/Saiga*, and *Equidae*, where sufficient sample sizes were available. While the overall deamidation trends across the full stratigraphic sequence remain broadly similar, distinct shifts were observed in layer 13 and layer 11.3, with elevated values for *pep1* and *pep2*, respectively. These layer-specific patterns could not have been identified using GA-q2e, which only targets pep2, nor by relying solely on the final MDS output that summarizes the overall deamidation rate for each sample. We focused on *pep1* and *pep2* as these peptides were the most well-defined in the spectra based on our quality control criteria, and their sequences remain conserved across most terrestrial mammal taxa. The deamidation trend across all layers of pep1 are very similar to both WLS-q2e on pep2 and GA-q2e, suggesting that pep1 may serve as an additional reliable marker for single-peptide deamidation analysis alongside the widely used pep2.

MDS returned insufficient data for heavier peptides, as most failed to pass the quality control filters. In some cases, false signals were generated when peptides with poorly defined isotopic envelopes as the case of *Equidae* grouping effect on pep4. When targeted peptides are absent due to poor preservation or instrumental bias, MDS may pick up isotopic peaks from unrelated peptides or even noise that happen to fall within the expected *m/z* range. Occasionally, these overlapping peaks mimic the isotopic distribution of a glutamine-containing peptide closely enough to pass the quality control steps, potentially introducing spurious data. For this situation, users can take advantage of the spectral inspection feature integrated into the preprocessing step, which allows them to revisit the original spectra and interpret the data more effectively, rather than passively accepting the model outputs as in GA-q2e.

### Archaeological implication

Overall, our MDS model not only reproduced the broad deamidation pattern observed with GA-q2e but also yielded more nuanced insight. In both approaches, a gradual decrease in deamidation is evident from layer 15 (and, where available, layer 17) through to layer 11.2, followed by a rapid increase beginning at layer 9.3. The shift between layers 9.3 and 9.2 is attributed to the influx of diagenetic agents, notably phosphates and humic acids derived from bat guano in the overlying Holocene layers (Shunkov et al., 2018, 2020). These acidic leachates infiltrated layer 9, substantially degrading bone preservation and promoting the formation of secondary phosphatic minerals such as taranakite and leucophosphite but did not penetrate beyond layer 11 (Brown et al., 2021c; Shunkov et al., 2020, 2018; Sokol et al., 2022), leaving the interval from layer 15 to layer 11 characterised by a steady temporal decline in deamidation with only localised diagenetic alterations (Morley et al., 2019; (Brown et al., 2021c; Shunkov et al., 2020, 2018; Sokol et al., 2022); Goldberg, personal communication).

Within this broader trend, peptide-specific patterns for pep1 and pep2 reveal additional complexities. Unexpectedly high deamidation was observed for pep2 in layer 11.3 and for pep1 in layer 13. While these results may simply reflect the small sample sizes for these layers, they could also suggest diagenetic processes that differentially affect the two peptides. The divergence between pep1 and pep2 trends in these layers may reflect highly localised processes acting on individual bones and different protein sequences. The phosphatic stratigraphy of the East Chamber provides some possible context. Layer 11.3 lies just beneath the strongly phosphatised zone of layer 9, where bat-guano-derived acidic leachates and phosphates are abundant (Sokol et al., 2022). Although the bulk geochemistry suggests that the phosphate front did not extend beyond layer 11, micro-scale seepage along fractures or porous pockets could have locally altered the chemistry in parts of layer 11.3, selectively accelerating the deamidation of more chemically susceptible peptides such as pep2. In contrast, layer 13 is well within the phosphate-free zone (layers 11.3–17; Sokol et al., 2022), yet (Zavala et al., 2021)) showed that even in these deeper units, sedDNA preservation varies markedly between micro-samples, likely due to patchy microenvironments influenced by organic inputs or localised pH fluctuations.

A further peptide-specific issue was detected for pep4 in *Equidae*, where a grouping effect was found to be an artefact caused by overlapping, unrelated peptides. This overlap lowered the estimated deamidation for pep4 in some spectra, possibly because preservation conditions had substantially reduced the targeted pep4 signal, allowing the irrelevant peptide to dominate (on the contrary, when the irrelevant peptide is significantly lower than the targeted peptide in terms of intensity it will be fully covered by the targeted peptide).

While the lack of taphonomic data for the Denisova assemblage analysed by ZooMS limits our ability to interpret these peptide-specific anomalies on individual bone samples, there is a growing trend to document taphonomic observations, where possible, for indeterminate bones in large-scale ZooMS studies (Lee-Michaelis et al., n.d.; Ruebens et al., 2024; Smith et al., 2024). Incorporating such information enables more detailed analyses of deamidation in relation to taphonomy and bone type (e.g., Welker et al., 2017; Ásmundsdóttir et al., 2025). Further application of MDS to datasets that include detailed morphological and taphonomic records would allow direct comparison of estimated deamidation rates with specific diagenetic variables, thereby greatly improving our understanding of potential links between deamidation and bone preservation.

### Clustering and batch effect

The estimation of deamidation is based on the isotopic peak distribution within the spectrum which in principle should not be affected by the batch effect. However the batch effect identified by the clustering analysis poses difficulty for large-scale ZooMS data analysis, as batch effects obscure meaningful biological or taphonomic patterns in the data. The sources of batch effect likely stem from multiple factors, including both batch (i.e., groupings of samples processed for collagen extraction) and plate (i.e., a plate of samples in the MALDI-TOF as a single run). Differences in instrument conditions, such as laser power settings or sensitivity losses over time, may contribute substantially to variability (Cai et al., 2023; Cuénod et al., 2023). Additionally, sample preparation protocols such as digestion time are often described as a range of time (from 12 to 18 hours or overnight), introducing further inconsistencies that could affect peptide recovery and deamidation estimates. Collectively, these technical factors generate noise that overwhelms potential subtle molecular differences between layers, limiting our ability to detect stratigraphic or taphonomic patterns. Addressing these sources of batch-related variation will be essential for future studies aiming to extract meaningful chronological or diagenetic information from large-scale ZooMS datasets.

We therefore recommend that researchers willing to do further quantitative analysis carefully consider batch effects when designing sampling strategies, for example by including consistent positive controls or deliberately mixing samples from different analytical units (such as layers) within the same batch to reduce batch-related bias. Nonetheless, such measures can only address batch effects within individual laboratories. To enable robust analysis of MALDI spectra across multiple laboratories, coordinated efforts will be required to mitigate batch effects, including the adoption of universal collagen standards for inter-laboratory calibration (Dekker et al., 2025).

## Conclusion

In this study, we introduce the MALDI Deamidation Score (MDS), a new statistical framework for estimating bone collagen deamidation from MALDI-TOF spectra, and demonstrate its application on the large faunal assemblage from Denisova Cave. Compared to the widely used q2e method, MDS integrates peptide-level estimates through weighted least squares and linear mixed-effects models, enabling the resolution of peptide-specific variation that could otherwise bias results. This formalisation provides a flexible framework that enables MDS to employ additional regression methods capable of better mitigating the stochastic noise characteristic of MALDI-TOF mass spectra. By accommodating user-defined peptide sets, triplicate runs, and partially failed spectra without manual filtering, MDS provides a transparent, scalable, and efficient solution for analysing large and taxonomically diverse ZooMS datasets.The application to Denisova Cave demonstrates how MDS can recover both broad deamidation trends and finer peptide-specific features, such as the elevated pep2 deamidation in layer 11.3 and pep1 in layer 13. These patterns, potentially linked to localised diagenetic microenvironments, illustrate the added interpretative value of peptide-level resolution. Such insights would have been obscured under aggregate deamidation estimates, underscoring the importance of integrating peptide-specific analysis into routine workflows.

A particular strength of MDS lies in its adaptability to ongoing advances in mass spectrometry technology. The framework’s flexibility in peptide selection and its robust statistical approach make it well-positioned for future applications as instrumentation continues to improve. As higher-resolution, lower-noise platforms such as MALDI-FTICR (Fourier transform-inductive coupled resonance) MS and TIMS-TOF MS become more widely available, spectral quality will improve, enabling reliable inclusion of peptides currently excluded due to poor isotopic patterns or low signal-to-noise ratios. The flexible structure of MDS will make the inclusion of additional peptides in future analyses straightforward, thereby improving the accuracy and reliability of deamidation-based reconstructions of taphonomy and site formation processes.

Realising the full potential of MDS will require pairing molecular-level properties (e.g., physicochemical) with detailed zooarchaeological and taphonomic records (e.g., bioclimatic and geochemical variables). Systematic contextual documentation will allow MDS outputs to be interpreted in relation to bone type, surface modifications, and diagenetic pathways.

MDS offers a flexible, scalable, and easy to interpret framework, thereby enabling us to measure glutamine deamidation in large-scale zooarchaeological datasets in a rapid and efficient manner. Even with limited metadata, our results show that MDS can yield valuable insights into diagenetic patterns. As measuring deamidation becomes routine in palaeoproteomic studies, MDS provides a powerful approach to explore biomolecular preservation, with the potential to be further strengthened through future integration with high-resolution mass spectrometry and detailed archaeological records.

## Data and code availability

The R package maldiMDS used to calculate the MALDI Deamidation Scores is available on GitHub (https://github.com/ismaRP/maldiMDS) and archived on Zenodo (https://doi.org/10.5281/zenodo.16938781). The scripts used to analyse the data and produce these results can be found on GitHub (https://github.com/ismaRP/MDS_Denisova) and archived on Zenodo (https://doi.org/10.5281/zenodo.16946802). It also contains the peptide table and metadata used. The data used in this study was obtained from the repositories https://doi.org/10.17632/bjvt47wb9c.1, https://doi.org/10.17632/wfsv6bt64w.1, https://doi.org/10.17632/h79h6t77tj.1 as they were archived by Brown et al. <u>(2021)</u>.

## Acknowledgement

We thank Paul Goldberg for his advice on the geoarchaeology of Denisova Cave. We thank Matthew Collins for engaging in discussions about deamidation and collagen preservation with us. Fei Yang was supported by The University of Tübingen’s Excellence Initiative. Ismael Rodríguez Palomo and Bharath Anila Bhuvanendran Nair worked on this study in their free time.

## Supplementary

**Figure S1.**
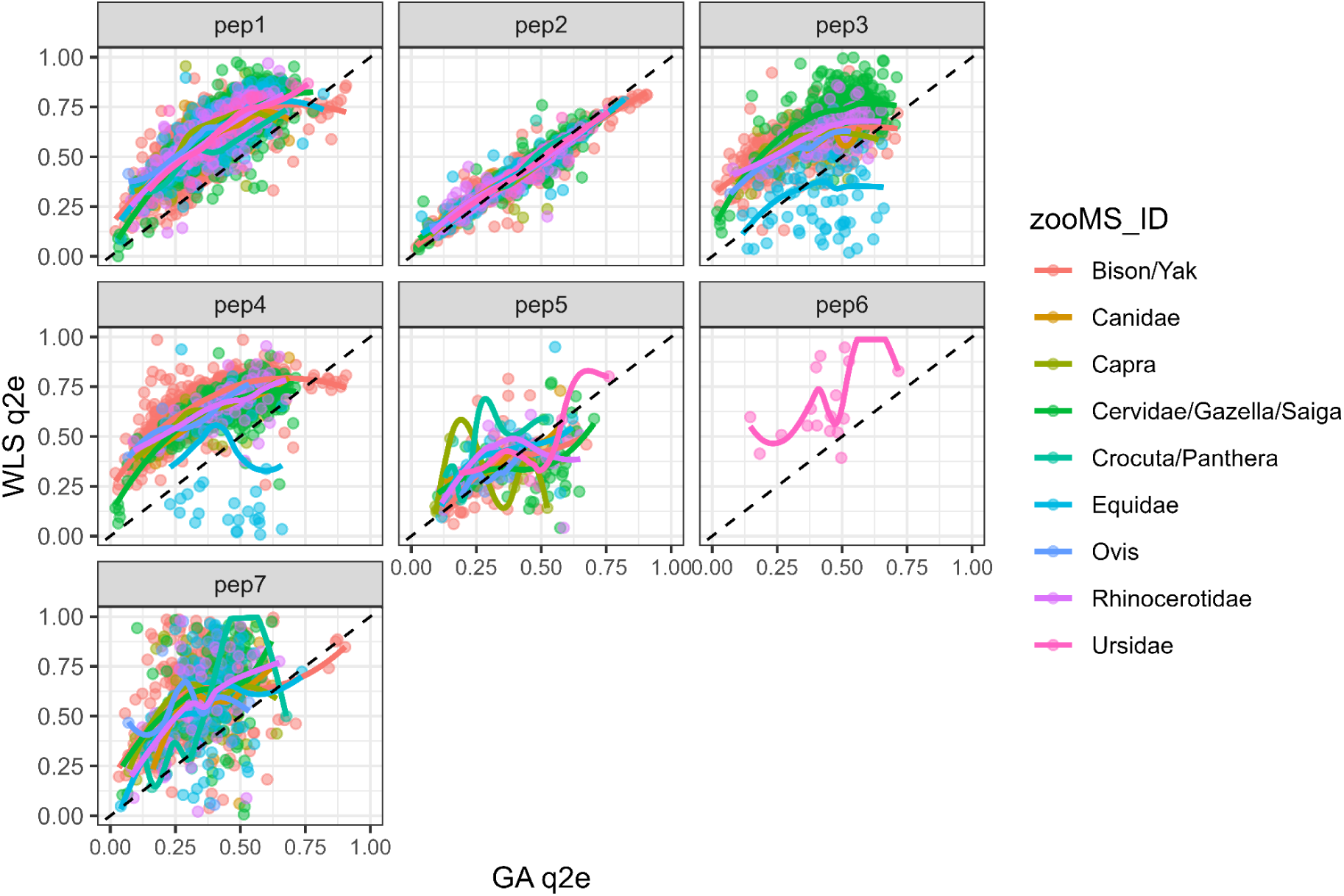
Comparison between WLS-q2e and GA-q2e estimates across peptides and taxa.

**Figure S2.**
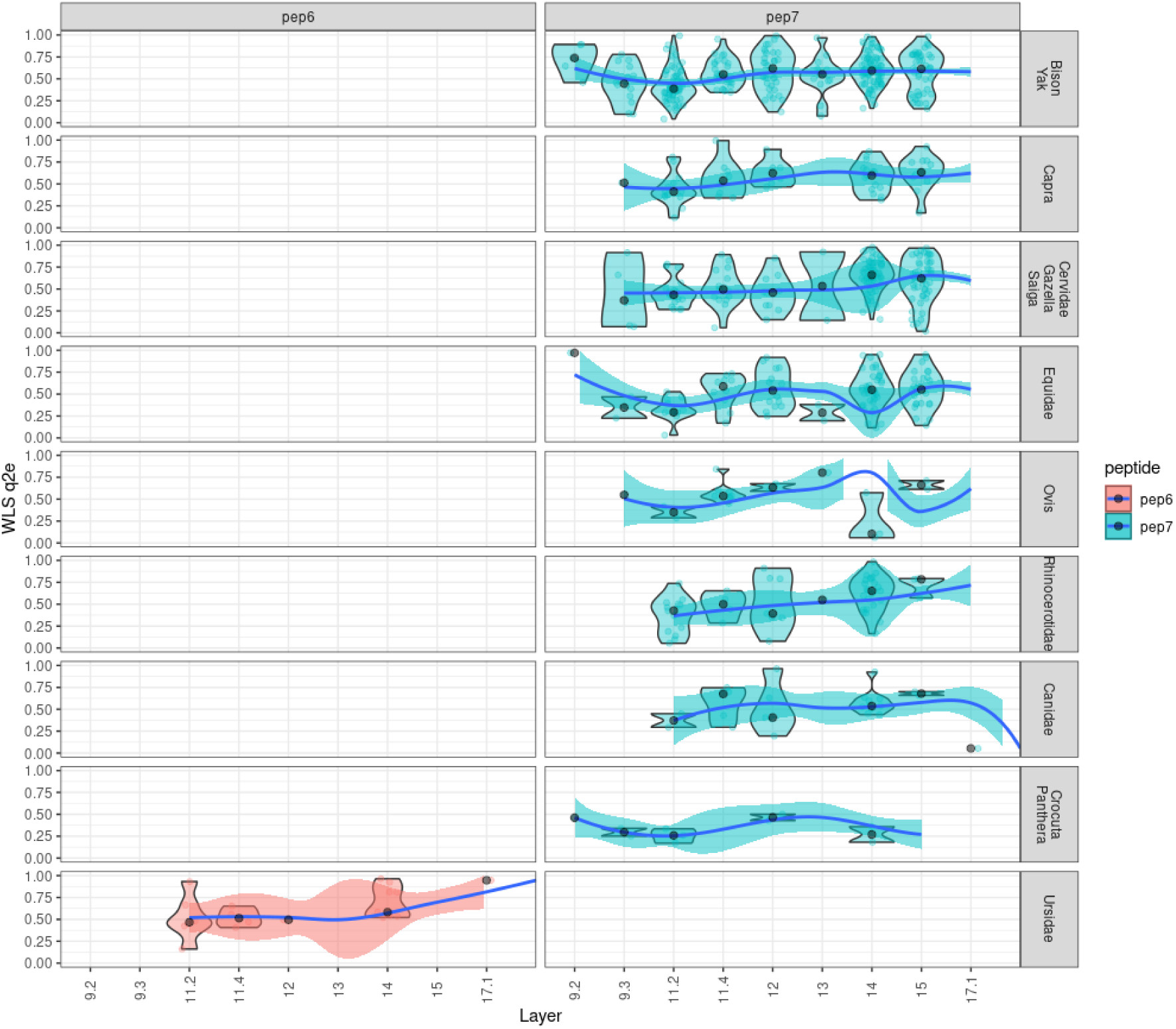
WLS-q2e patterns across stratigraphic layers for pep6 and pep7.

**Figure S3.**
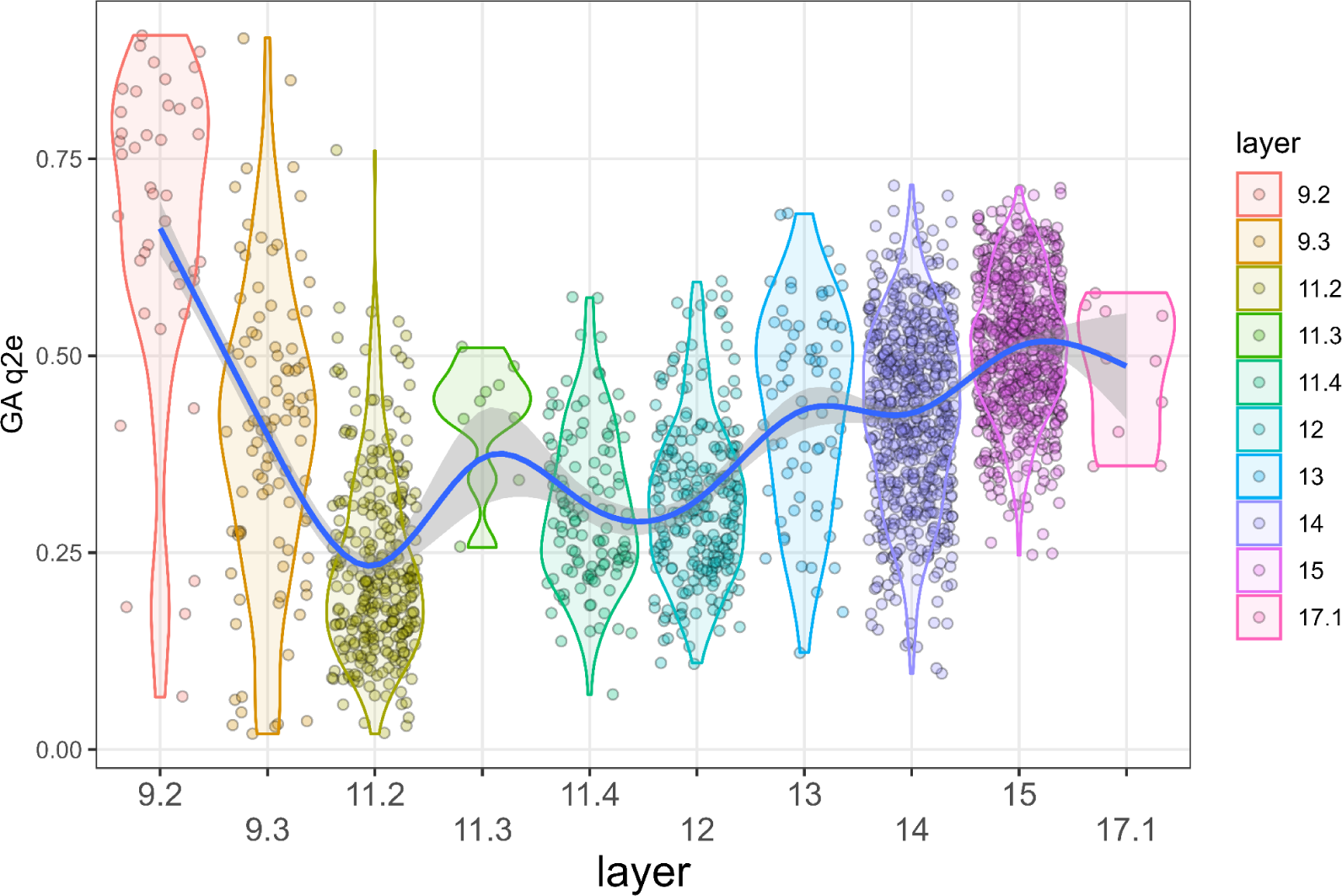
Diachronic trends of GA-q2e across layers.

